# Synchronized Temporal-spatial Analysis via Microscopy and Phospho-proteomics (STAMP) of Quiescence

**DOI:** 10.1101/2024.06.24.600542

**Authors:** Mohammad Ovais Azizzanjani, Rachel E. Turn, Anushweta Asthana, Karen Y. Linde-Garelli, Lucy Artemis Xu, Leilani E. Labrie, Mohammadamin Mobedi, Peter K. Jackson

## Abstract

Coordinated cell cycle regulation is essential for homeostasis, with most cells in the body residing in quiescence (G_0_). Many pathologies arise due to disruptions in tissue-specific G_0_, yet little is known about the temporal-spatial mechanisms that establish G_0_ and its signaling hub, primary cilia. Mechanistic insight is limited by asynchronous model systems and failure to connect context-specific, transient mechanisms to function. To address this gap, we developed STAMP (**S**ynchronized **T**emporal-spatial **A**nalysis via **M**icroscopy and **P**hospho-proteomics) to track changes in cellular landscape occurring throughout G_0_ transition and ciliogenesis. For the first time, we synchronized ciliogenesis and G_0_ transition in two cell models and combined microscopy with phospho-proteomics to order signals for further targeted analyses. We propose that STAMP is broadly applicable for studying temporal-spatial signaling in many biological contexts. The findings revealed through STAMP provide critical insight into healthy cellular functions often disrupted in pathologies, paving the way for targeted therapeutics.

**TEASER:** STAMP of signaling in quiescent cells unravels transient phosphorylations in cellular functions.

## INTRODUCTION

Fundamental cellular processes are driven by highly coordinated, temporal-spatial regulation of cell signaling networks. While proliferation and differentiation require major changes in transcriptional programs, most cells in adult tissues are not proliferating either as post-mitotic or periodically regenerating cells. These non-proliferating cells are often engaged in physiological signaling and often use rapid, protein-driven regulatory mechanisms. Post-translational modifications (PTMs), in particular, allow for rapid and readily reversible cellular responses to environmental cues, as these transient and frequently reversible modifications can alter protein structure, stability, interactions, and localization. The transient nature of PTMs (including phosphorylations, the most widely studied of PTMs) and protein localization underscores the dynamic and reversible nature of cellular regulation. Disrupting these critical PTMs and transient, context-specific cellular functions can lead to severe pathologies, highlighting the urgent need to dissect these mechanisms. Yet, because many previous studies have relied upon cancer cell lines as model systems for signaling, we have limited insight into the signaling landscape of healthy, quiescent (G_0_) cells, the most prevalent cell cycle status in human tissues(*2*). Identifying transient phosphorylations that induce or later maintain quiescence (G_0_) in noncancerous cells is crucial not only for advancing cell biology but also for human health, as the majority of pathologies arise from a failure of cells to either enter quiescence or to perform essential fated, quiescent cell functions. This includes cancers(*3–5*), neural aging(*6*), and chronic metabolic diseases(*7*). Mapping the temporal-spatial signaling landscape in healthy, quiescent cells is complicated, though, by the dearth of quiescent program markers and the lack of tools to track changes in PTMs and protein function in time and space.

One of the few widely accepted hallmarks of quiescence in most cell types is the primary cilium, an organelle that protrudes into the extracellular space and acts as a pivotal signaling hub(*8*). This tiny organelle is composed of the mother centriole, or basal body, that projects a microtubule-based axoneme encased in a ciliary membrane that is contiguous with the plasma membrane. The primary cilium houses specific receptors that sense extracellular stimuli and transduce those signals to propagate discrete cellular responses. Disruption of these signals drives a number of severe pathologies, including multi-system developmental disorders called ciliopathies, as well as tissue-specific diseases including blindness, kidney disorders, diabetes, obesity, Parkinson’s disease, and cancers(*9–18*).

A significant impediment in studying G_0_ transition and how cilia are formed and function in G_0_ is the inability to detect transient, context-specific signals in time and space. For example, the standard method to induce ciliogenesis in cultured cells involves seeding at a high density (80-90%) followed by serum starvation for 24-72hrs. However, the lack of synchrony in G_0_ entry and asynchronous initiation of ciliogenesis suppresses consistent observation of transient changes to below the limits of detection by either imaging or proteomics; this results in the loss of the temporal dimension of both cellular and molecular events. To tackle this challenge, we first developed an improved protocol for synchronous cell cycle progression into G_0_, focusing very specific events into narrow time windows. We then leveraged the power of state-of-the-art mass spectrometry (MS) and fixed or live-cell imaging to track the temporal-spatial network of proteins and their PTMs in differential signaling contexts. By generating much more homogeneous cell populations, we can effectively correlate changes in cellular processes via microscopy with changes in specific transient PTMs identified by mass spectrometry. Wedding both approaches enables testable, targeted hypotheses informing mechanisms. Our approach for teasing apart subcellular temporal-spatial PTM-driven signaling is termed **STAMP**: **<u>S</u>**ynchronized **<u>T</u>**emporal-spatial **<u>A</u>**nalysis via **<u>M</u>**icroscopy and **<u>P</u>**hospho-proteomics. Below, we provide a detailed methodology for applying STAMP to tracking temporal phosphorylations and their correlated effects on the localization and function of these proteins on cytokinesis, nuclear pore complex (NPC) formation, Golgi reorganization, autophagy, centrosomal maturation, translation, and ciliogenesis from mitosis to G_0_ progression in non-cancerous cells. We focus here on our analyses of G_0_ and ciliogenesis to demonstrate the value of the methodology, though note that it can be used to dissect and identify key components in many other cellular processes.

## RESULTS

### Premise

Our goal was to develop a method to allow detailed microscopy-based cellular analyses of distinct stages in the transition from mitosis to G_0_ commitment, including but not limited to mitosis, cytokinesis, ciliogenesis, translational control, and vesicular traffic. The homogeneous appearance of specific cellular events would be expected to reflect the underlying molecular state. This expectation of molecular synchrony can be confirmed by demonstrating that the phosphoproteome profile of cells are similarly homogenous in each time window, thus linking timed cell biological events to phosphorylations that define each temporal window.

To achieve this goal, we developed a cell synchronization protocol that yielded a pure population of metaphase cells and released them into serum-free medium to track the time-dependent events driving G_0_ progression, followed by multi-dimensional analyses with high temporal resolution over 24hrs, the time required for ciliation and completion of other synchronous, stepwise G_0_ events. The combination of fixed and live cell imaging with state-of-the-art phospho-proteomics (Fig.1A) allowed us to track the detailed progression of cellular events (Fig.2, Fig.3) and their associated phosphorylation states (Fig.4). By doing so, we observed strong correlations that suggest testable hypotheses regarding the roles of specific proteins and their phosphorylations (often transient or rare and previously undetected) in regulating function and localization. As our model system, we primarily used hTERT-immortalized human retinal pigment epithelial cells (hTERT-RPE-1) as they are: 1) human, non-cancerous, adherent, mononucleated cells with a very stable karyotype; 2) a commonly used model system for studies of ciliogenesis and cell cycle, being highly-regulated by the Rb tumor suppressor pathway(*19*); and 3) readily tractable for many cellular perturbations and signaling assays. We have also successfully performed STAMP of quiescence and ciliogenesis in Mouse Embryonic Fibroblasts (MEFs), as described below, highlighting the versatility of our approach.

**Fig.1:**
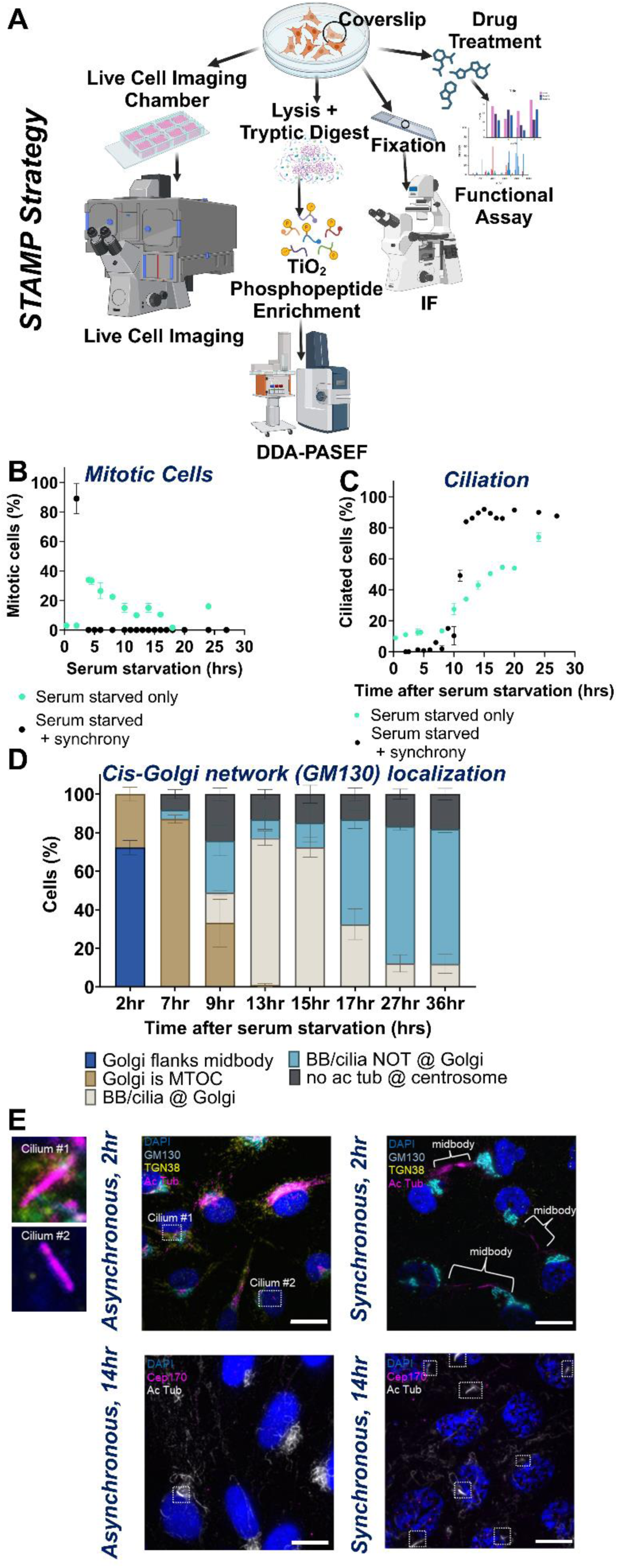
Rationale for the development of STAMP to dissect cell signaling. (A) Schematic representing the various applications for synchronized cell populations, summarizing our approach for STAMP as a way of deciphering temporal-spatial mechanisms via live and fixed cell imaging as well as detailed mass spectrometry (in this case, phospho-proteomics). Created in BioRender. Turn, R. (2025) https://BioRender.com/v05m172. (B-D) Quantification of mitotic indices, ciliation rate, Golgi morphology, and Golgi versus centrosomal MTOC formation in STAMP-synchronized versus serum starved only cell populations, based on IF. These experiments were performed in biological replicate, with N=100 cells per replicate. Error bars indicate SEM. (E) Example IF images of cells that were binned based on ciliation, Golgi morphology, MTOCs, or mitotic indices. For the 2hr timepoint, cells were stained for DAPI (blue), TGN38 (yellow), GM130 (cyan), and acetylated tubulin (magenta). For the 14hr time point, markers used were DAPI (blue) to mark nuclei, acetylated tubulin (white) to mark mitotic microtubules and primary cilia, and CEP170 to mark the basal body. Images were collected using widefield microscopy, 60x magnification, z-projection. Scale bars= 10μm.

**Fig.2.**
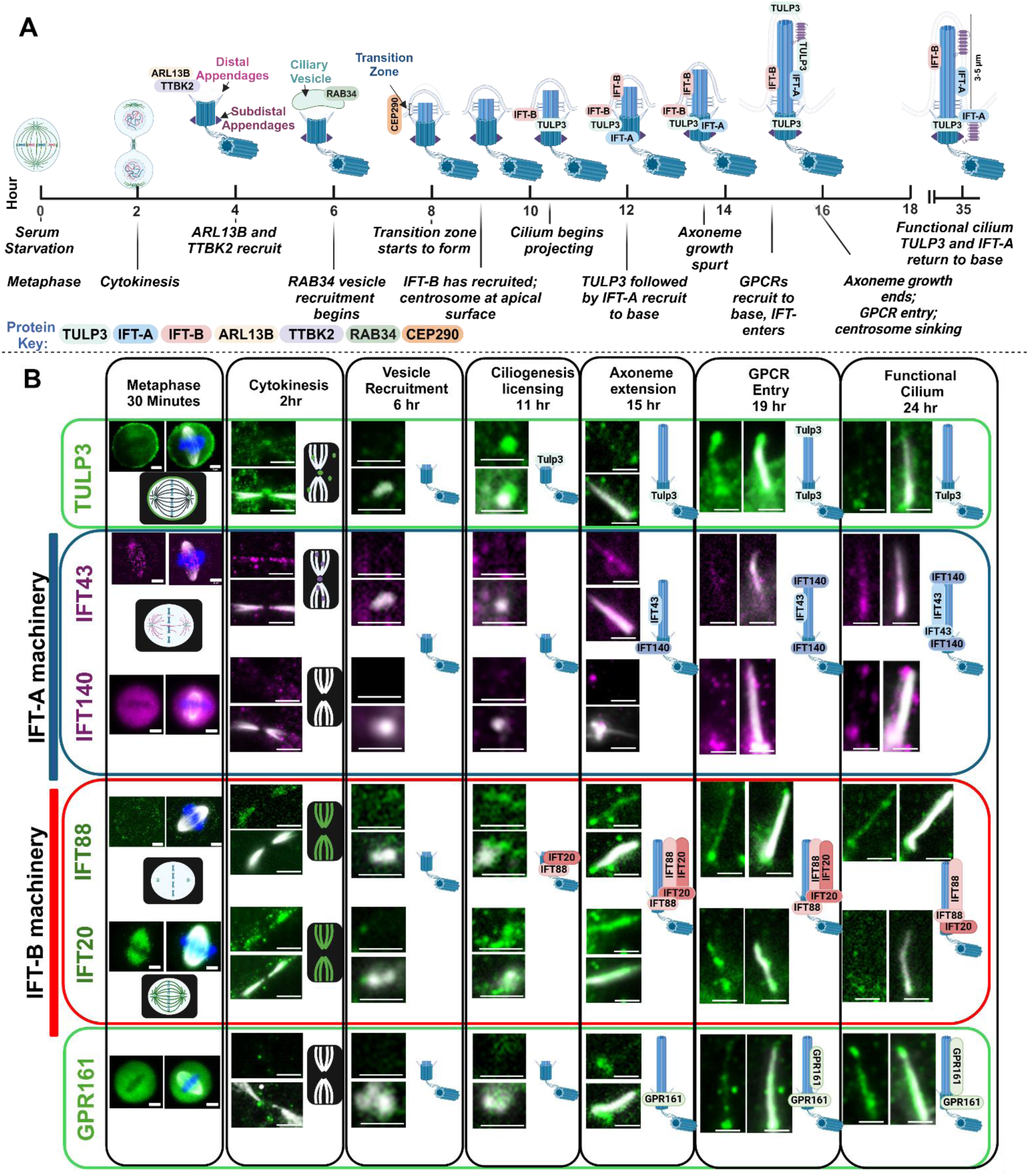
Developing a temporal-spatial map of quiescence via fixed cell imaging. (A) Temporal-spatial map of the stages of exit from mitosis and cytokinesis and progression into G_0_, as marked by ciliogenesis. This map was based on cells plated in serum-free medium on fibronectin coated plates, collecting time points approximately every hour over a 24hr period. This map summarizes the findings from compiled IF and live cell imaging data, collected from over multiple iterations of the STAMP protocol. The results were highly reproducible, with the temporal localization of key markers being consistent (within a 1-2hr margin) between experimental runs. (B) Representative immunofluorescence images of key stages of IFT-A (IFT140 and IFT43) IFT-B (IFT20 and IFT88), TULP3, and GPR161 recruitment and localization to the cilia and basal body are shown below the map (widefield microscopy, 60x magnification, z-projection). Markers for each of the images are labeled with the respective name and color of the IF marker used. Created in BioRender. Turn, R. (2025) https://BioRender.com/y11n181. Scale bars for centrosomes and cilia = 2μm; Scale bars for midbodies and metaphase cells = 5μm.

**Fig.3.**
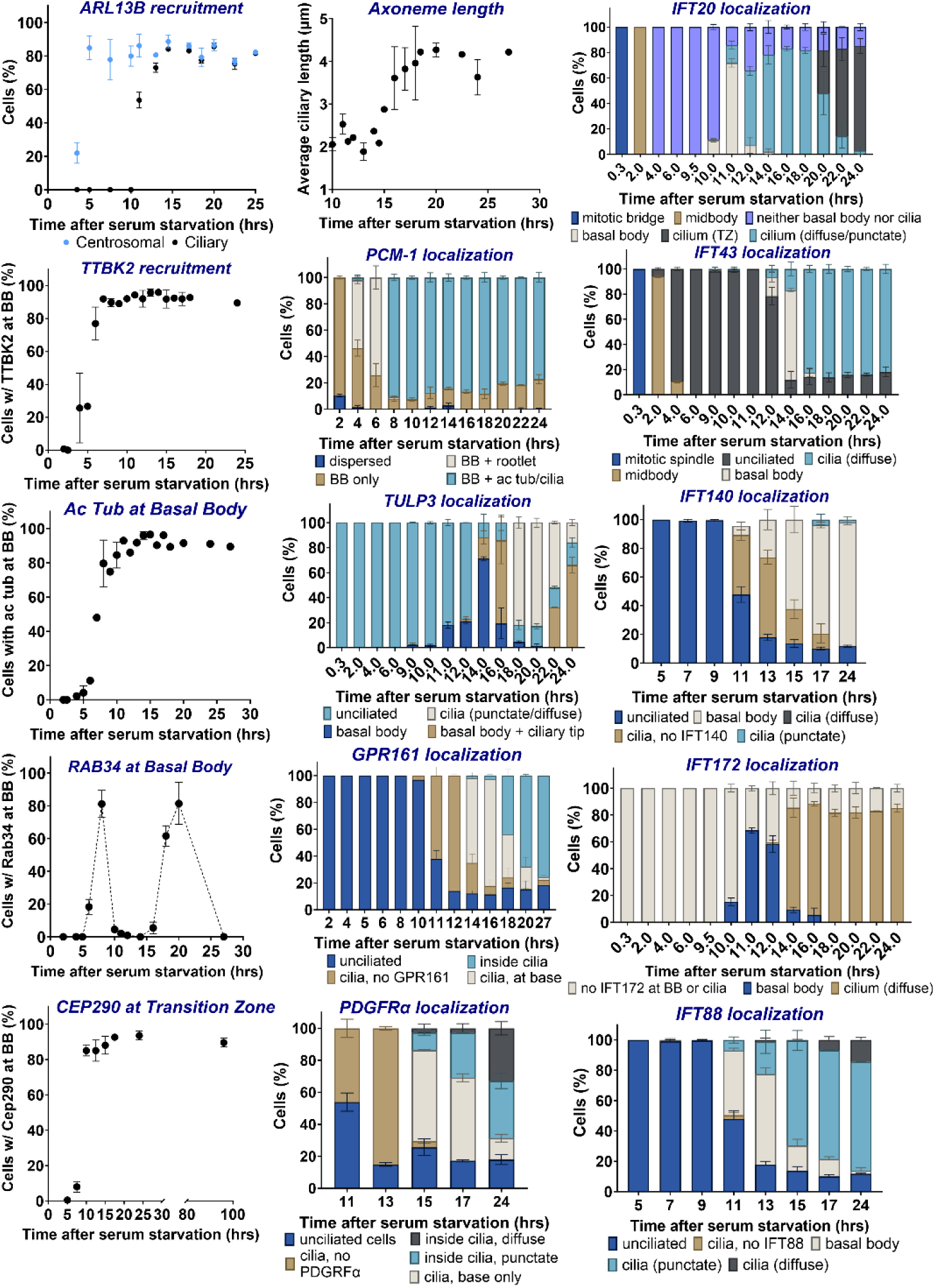
Localization of different markers of cell cycle/ciliogenesis/key cellular compartment morphologies are quantified to establish the temporal-spatial landscape of G_0_ progression. Experiments were performed in replicate and quantified over a range of time points after plating onto fibronectin coated coverslips. N= 100 cells per replicate, error bars= SEM.

**Fig.4.**
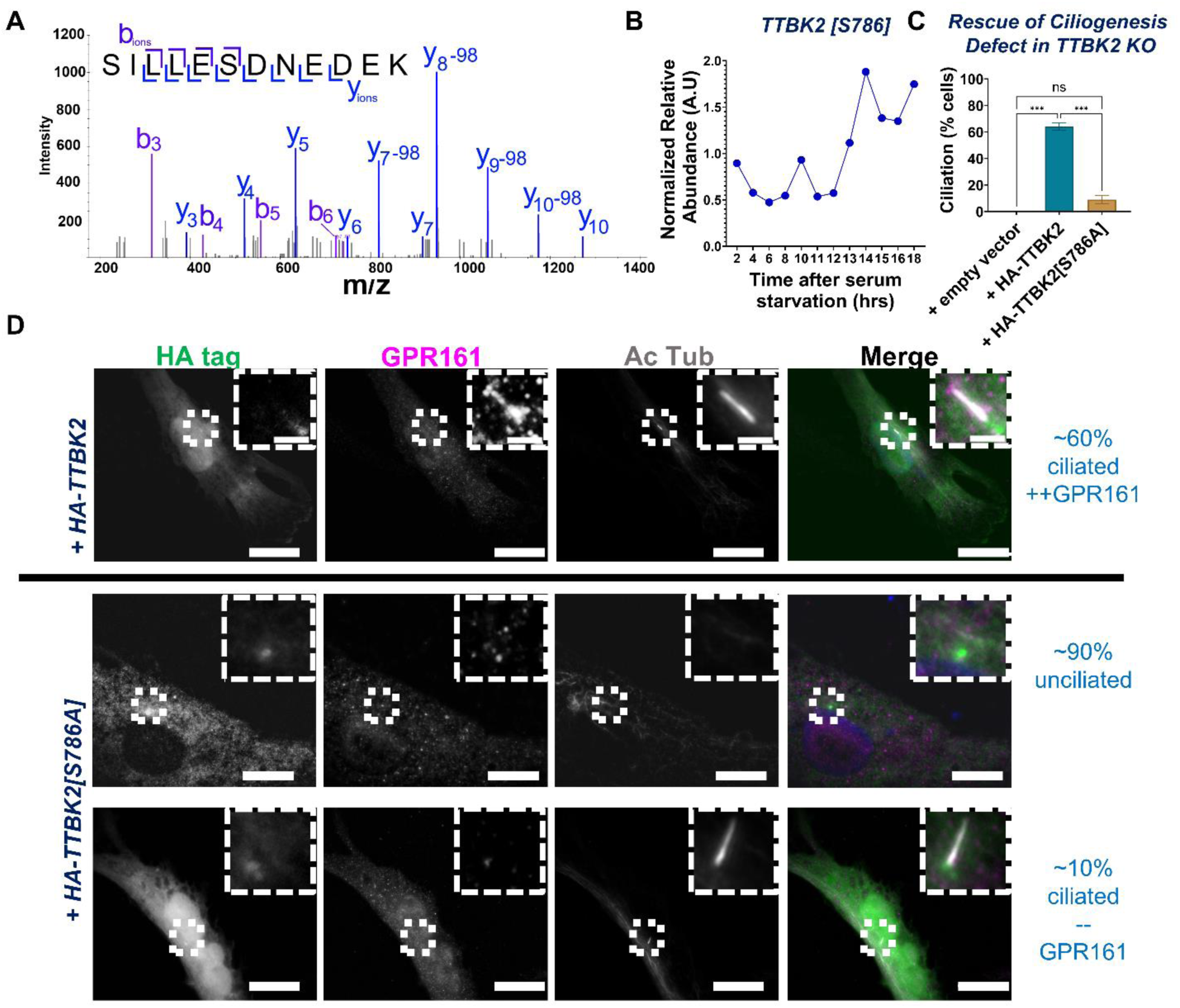
TTBK2[S786] phosphorylation is a critical regulator of late-stage ciliogenesis. (A) MS2-CAD scan of TTBK2[S786] identified via STAMP phospho-proteomics. These spectra were used to confirm which residue is modified for this phosphopeptide. (B) Quantification of TTBK2[S786] phosphorylation levels over time via mass spectrometry revealed a spike in abundance during late ciliogenesis stages, specifically axonemogenesis and GPCR recruitment windows. (C) Quantification of the percentage of ciliated cells shows that HA-TTBK2 rescues ciliogenesis (∼60% ciliation), while HA-TTBK2[S786A] fails to do so (∼10% ciliation). Error bars represent SEM; statistical significance was determined using One-Way ANOVA. Scoring was performed in replicate, with N=100 cells per replicate. (D) Representative images of TTBK2 KO RPE cells transiently transfected with empty vector, HA-TTBK2, or HA-TTBK2[S786A] (phospho-defective mutant), followed by serum starvation and immunofluorescence (IF) staining. Antibodies against HA (to detect transfected cells), acetylated tubulin (axoneme marker), and GPR161 (to assess GPCR recruitment) were used. Scale bar = 10μm.

### Generation of synchronized cells

We developed a rigorous strategy to synchronize the transition to quiescence, using ciliogenesis as the key marker of progression through G_0_. The established approach for inducing G_0_ and ciliation is serum starvation (0-0.5% fetal bovine serum) of cells near confluence (80-90% density). Primary cilia can be identified via immunofluorescence (IF) using antibodies targeting acetylated tubulin to mark the axoneme, ARL13B to mark the ciliary membrane, and Cep170 or centrin to mark the basal body(*20–25*). When asynchronous cell populations grown on fibronectin-coated coverslips were switched to serum-free medium, we observed continuous but asynchronous progression of ciliary assembly, with cilia forming as early as 2hrs and gradually increasing in number to 70-90% ciliated cells over a 24hr window (Fig.1B-C). Note that this standard approach of tracking ciliogenesis only accounts for axoneme extension and lacks data on the timing of earlier licensing events or functionality in receptor recruitment. This asynchronous progression of cells undergoing ciliogenesis does not allow tracking or ordering of transient cellular events or linking them to transient changes in PTMs. Therefore, we developed a validated, multi-step synchronization protocol to achieve much improved time-resolved analysis of the specific steps progressing through G_0_, including key steps needed to establish ciliogenesis.

After testing multiple protocols, drug concentrations, and incubation times to maximize cell viability, synchrony, and yield, we developed the following protocol for synchronizing G_0_ entry in RPE cells. Cells were seeded at 10% confluence and then grown in standard growth medium with 10% serum for 24hrs, at which point the CDK4 cyclin-dependent kinase inhibitor palbociclib (1μg/mL) was added to arrest cells in late G_1_ (Fig.S1A). After 18hrs of G_1_ synchronization, the drug was washed out, and the cells were allowed to recover for 8hrs. At this point, the microtubule inhibitor nocodazole (50ng/mL) was added to synchronize cells at metaphase. After 12hrs, we performed mitotic shake-off to isolate metaphase-synchronized cells, achieving ∼20-30% yield of total cell input. Based on immunostaining, this triple synchronization methodology (double block and shake-off) yields nearly 100% metaphase-synchronized cells (Fig.1B). We then filtered for single cells using a cell strainer and incubated in standard growth medium with serum for 20min at 37°C before plating. This allowed microtubules to recover, facilitating improved adhesion and earlier, more synchronous ciliogenesis (Fig.S1B) than serum withdrawal alone. Cells were then plated onto fibronectin-coated coverslips and/or plates in serum-free medium to induce G_0_ progression(*26, 27*). By monitoring multiple events, we confirmed a uniquely high degree of synchronization in mitosis and cilia formation, with essentially all cells completing cytokinesis by 2hr and initiating cilia projection within a 2-hour window approximately 10-12hrs after seeding (Fig.1C-E). Previous live cell imaging studies of axonemogenesis in untreated RPE cells(*22*) align with our findings in synchronized cells following drug treatments, with both showing that it takes approximately ∼100 minutes for a cilium to reach its full length from the start of extension(*28*).

### Temporal changes in protein localization reveal stages of quiescent transition

With evidence that the protocol achieved a high degree of synchronization, we fixed cells for IF every 1 or 2 hours for 24 hours following recovery from mitotic shake-off to document cellular changes, using previously well-characterized markers. These included: CEP164 and TTBK2 as markers of centrosome maturation; TGN38 and GM130 for Golgi morphology; RAB34 to track vesicle transport to the centrosome; CEP290 to label the transition zones; GPR161, PDGRFα, and ARL13B as representative ciliary membrane proteins; acetylated tubulin to both identify axonemes and determine cilia lengths; intraflagellar transport (IFT) components IFT88, IFT20, and IFT172 to identify different populations of the IFT-B complex at the centriole or in the cilium; and IFT43 and IFT140 to mark the IFT-A complex, along with TULP3 as a critical regulator of IFT-A(*21*). Recruitment/localization of these markers was tracked across multiple replicates and quantified, highlighting the reproducibility of our approach, as shown in Figs.2-3.

This allowed us to build a temporal map of the steps involved in G_0_ entry and ciliogenesis, summarized in Fig.2A(*20–22, 29–31*). These include: **1)** completion of cytokinesis by ∼2hr; **2)** recruitment of ARL13B and TTBK2 to the basal body within ∼3-4hr; **3)** RAB34 containing ciliary vesicle recruitment to the centrosome at ∼6hr, with its loss by ∼10hr; **4)** transition zone formation (marked by Cep290 and acetylated tubulin) at ∼9hr; **5)** IFT-B recruitment to the basal body (marked by IFT88, IFT20, IFT172) and ciliation (marked by acetylated tubulin and IFT-B train entry) at ∼10-11hr; **6)** TULP3 recruitment at ∼11hr followed by IFT-A (IFT140, IFT43) recruitment at ∼13hr; **7)** axonemogenesis occurring over a 4hr window from ∼13-17hr; **8)** GPCR entry into cilia at ∼18hr-20hr. Note that some of these key signals have transient localizations inside the primary cilia and/or at the basal body(*32, 33*). Notably, RAB34 has two waves of localization to the basal body: first during ciliary vesicle recruitment (a key intermediate of ciliary assembly) (7-10hr) and second during GPCR recruitment (17-20hr). This is consistent with previous work suggesting multiple functions for Rab34 in primary cilia(*33*). TULP3 also demonstrates transient localization along the length of the primary cilium between 13-20hrs, consistent with its roles in ciliary receptor entry(*21*).

These findings highlight the value of using synchronized model systems to track temporal-spatial cellular events, as such transient signals and their corresponding molecular mechanisms may be missed in other contexts. For example, this is the first evidence, to our knowledge, that IFT-B is recruited to basal bodies *before* IFT-A and that cilia appear to need to be fully formed before GPCR entry (Fig.2-3). Furthermore, our data reveal key differences between individual components of the IFT-A and IFT-B complexes: that the individual subunits themselves have discrete temporal localization patterns. In the case of IFTB components, IFT172 and IFT88 have similar temporal-spatial distribution: recruiting early to the basal body, entering during axonemogenesis, and remaining diffuse in the cilium throughout the time course. On the other hand, IFT20 only localizes transiently inside the cilium, returning to the base of the cilium by the end of the GPCR entry window (∼20-22hr). We also observe IFT20 localization to Golgi beginning at ∼6-7hr, consistent with previous reports in the literature of IFT20 regulating the traffic of ciliary membrane proteins(*34*). When we track IFT-A components, we observe that IFT43, like IFT140, recruits to the basal body later (∼13-14hr). Endogenous staining of IFT43 differs from IFT140, though, as we observe diffuse staining of IFT43 that persists even after GPCR recruitment. It is also interesting to note that IFT43, like IFT-B components, localizes to the mitotic spindle and the cytokinetic bridge (Fig.2). Altogether, these findings reveal critical details concerning the stages of ciliogenesis and lay the groundwork for probing how and when defects in specific components alter these processes.

We also mapped time-dependent changes in other key cellular compartments, including trans-Golgi (TGN38) and cis-Golgi (GM130) network reassembly (Fig.1D,E 2hr window). The Golgi dynamically reorganizes in relation to the centrosome and basal body over time. After cytokinesis, the Golgi (GM130) serves as the primary MTOC (MicroTubule Organizing Center) at around ∼7hr. By ∼9hr, the acetylated tubulin-positive microtubules no longer show strong colocalization at the Golgi and instead begin to localize to the centrosome. Between 13-15hr, the basal body and growing primary cilium are proximal with Golgi. By ∼17hr after release (which is post-axonemogenesis), the cilium shows reduced incidence of Golgi proximity (Fig.1D). We also note that, as cells transition from mitosis further into the G_0_ program, the number and intensity of TGN38-positive TGN compartments reduces over time (Fig.1E). Future work can be done to establish the functionality of these compartments in G_0_ and how they respond to different stimuli.

To determine if STAMP can be applied to other types of cells, we repeated these studies using SV40 immortalized MEFs (Fig.S2). Although different cell types vary in drug sensitivities, cell doubling times, or extent of ciliation, we found that synchronization of MEFs at metaphase produced a slightly higher yield than RPE cells. Staining of the same ciliary markers in both MEFs and RPEs revealed conservation of the order of events but different kinetics for ciliogenesis. MEFs attached to the fibronectin-coated coverslips/plate much faster than RPE cells, already flattening on the plate at 20min post-seeding, while RPE cells are still rounded and in early cytokinesis. Ciliation also occurred at earlier time points in MEFs (∼7hr) compared to RPE cells (∼12hr). Together, these data reflect the general applicability of this synchronization strategy to diverse cell lines, though highlighting the need for optimization to map kinetics. Such differences in kinetics between cell types might be exploited in later studies to address the conservation or divergence of pathways and their components.

### Phospho-proteomics reveals transient phosphorylations at distinct time points from mitosis to G_0_ entry

The high degree of cell synchronization achieved with our protocol allowed us to identify and further analyze transient events that otherwise are undetectable in a more heterogeneous cell population. Transient changes in PTMs drive coordinated cellular functions, so we employed phospho-proteomics to monitor phosphorylations at each time point shown in Fig.2A. We simultaneously harvested samples for IF and phospho-proteomics by seeding a coverslip in the same 10cm plate used to harvest cells for phospho-proteomics (Fig.1A). We employed data-dependent acquisition (DDA) in Parallel Accumulation-Serial Fragmentation (PASEF) mode to identify phosphorylation sites(*35*). This approach involved fragmenting ions from the MS1 signal based on their relative abundance every 100ms, which may result in the loss of low-abundance features due to instrument sensitivity and sample complexity. However, DDA is essential for obtaining high quality MS2 scans for confirming phosphorylation sites. While advanced instruments continually improve detection limits every few years(*36*), in heterogeneous cell populations, uncontrolled variability often obscures relative phosphorylation changes, limiting the detection of transient phosphorylation events. STAMP phospho-proteomics addresses this limitation by synchronizing cells to generate a much more homogeneous cell population, enriching phosphorylation events that might otherwise be diluted in heterogeneous samples. This enrichment effectively reduces sample complexity and enhances sensitivity, allowing us to detect transient phosphorylation events with improved robustness (Fig.S1C).

We posited that STAMP may serve as a powerful tool for predicting new phosphoregulatory steps in the ciliogenesis and G_0_ programs, even in the case of low abundance phosphosignatures. As proof of principle, we looked through our phospho-proteomics data to identify candidate regulators of the ciliogenesis program. Among the many candidates we identified, we were struck by one phosphorylation: TTBK2[S786] (Fig.4A).

TTBK2 is a critical kinase known for licensing ciliogenesis, phosphorylating key substrates necessary for ciliogenesis(*29, 37*). Loss of TTBK2 prevents ciliogenesis, and defective TTBK2 leads to severe pathologies including Spinocerebellar Ataxia and Alzheimer’s Disease(*29, 38–40*). This low-abundance protein is often difficult to detect via mass spectrometry, much less phospho-proteomics. Via STAMP, though, we could detect TTBK2[S786] and track its abundance over time. This phosphorylation is in high abundance late in the ciliogenesis program, spiking specifically during the axonemogenesis and GPCR recruitment windows (Fig.4B). We found this trend especially surprising, as most of the known functions for TTBK2 are specifically in relation to driving the early ciliogenesis program.

We decided to test whether TTBK2[S786] regulates ciliogenesis by comparing its ability to rescue ciliation in TTBK2 KO RPE cells. We transiently transfected empty vector, HA-TTBK2, and HA-TTBK2[S786A] (phospho-defective mutant) in TTBK2 KO RPE cells, serum starved the cells for 24hrs, and quantified the number of ciliated cells via IF (Fig.4C-D). We used antibodies against HA (to detect transfected cells), acetylated tubulin (to identify the axoneme of ciliated cells), and GPR161, a key ciliary GPCR, to determine whether GPCR recruitment is defective. We observed that both WT and TTBK2[S786A] localize to basal bodies, suggesting that this mutation does not affect TTBK2 localization. However, TTBK2[S786A] failed to rescue ciliogenesis. While transient expression of HA-TTBK2 in TTBK2 KO background leads to ∼60% ciliated cells, HA-TTBK2[S786A] transient expression only leads to ∼10% ciliation (Fig.4C). Furthermore, the few cilia that did form upon expression of HA-TTBK2[S786A] failed to recruit GPR161. Altogether, these data point to a critical role for phosphorylation of TTBK2[S786] in driving late-stage ciliogenesis.

As another example, STAMP phospho-proteomics successfully identified transient phosphorylation events on CEP131, capturing both frequently and infrequently detected sites based on data in PhosphoSitePlus®(*1*) (Fig.S3). CEP131 is a centriolar satellite protein which regulates the recruitment of key ciliogenesis factors to the basal body(*41*). The CEP131 sites that we identified exhibit an increase in over an order of magnitude in intensity during G_0_ progression, demonstrating the robustness of our quantification and supporting the detection of low-abundance phosphorylation events (Fig.S3). Specifically, proline-directed phosphorylation sites of CEP131 are enriched during cell division, whereas N-terminal sites, such as [S47] and [S78], are phosphorylated during vesicle recruitment to the basal body but diminish afterward. To validate the functional relevance of our findings, we reference previous studies demonstrating that CEP131[S78] phosphorylation is essential for centriolar satellite integrity(*42*). In those studies, depletion of CEP131 resulted in increased dispersion of centriolar satellites, which could be rescued by wild-type and phosphomimetic [S78D] CEP131 but not by the non-phosphorylatable [S78A] mutant(*42, 43*). These findings provide strong evidence that our phospho-proteomics data align with known biological functions and confirm the significance of phosphorylation events in specific time windows and cell cycle progression. Future work will include targeted mutagenesis to further validate additional phosphorylation sites identified through our approach and to examine the function of centriolar satellites in vesicle recruitment to primary cilia.

STAMP phospho-proteomics allowed us to optimally correlate cellular changes to both transient and stable phosphorylations in RPE cells. We observed >9700 phosphorylation events (including several double and triple phosphorylations) using TiO_2_ enrichment and the timsTOF HT mass spectrometer. We provide an example of how we analyzed phospho-signatures in Fig.S4, looking at 5 different phosphosites on the ULK1 S/T-protein kinase (a key regulator of autophagy) and their relative fold change in normalized intensity over time. ULK1[S450] is an example of a very stable phosphorylation 8 hours post-mitosis. We identified several such phosphorylations that remained constant throughout the time course, suggesting that their role was for general cellular functions (*e.g*, proteostasis, metabolic adaptation). In contrast, other phosphorylations showed dynamic changes in relative abundance that temporally correlated with specific stages of G_0_. For instance, the double phosphorylated peptide (NLQ<u>pS</u>PTQFQ<u>pT</u>PR), ULK1[S450-S456], is four-fold higher in relative abundance during mitosis, suggesting the importance of different ULK1 phospho-proteoforms during cell cycle progression and G_0_. For phospho-signatures of interest, we searched the literature to identify known functions. Though all these sites were reported in PhosphoSitePlus, their functions are unknown except in the case of ULK1[S469], which is reported as promoting the ubiquitin-dependent degradation of ULK1. Interestingly, ULK1[S469] appears and gradually increases in abundance starting at around 4-5hr, suggesting its role in degradation of the mitotic ULK1 pool. Overall, homogeneous cell populations allowed us to track context-specific, transient phosphorylations even on low-abundance proteins, many of which were identified less frequently or were missing in other published phosphoproteomic screens(*1*). Fewer than 10% of the phosphorylations that we identified to date have an annotated, known functionality based on PhosphoSitePlus, and many of the reported functions for the “known” sites are inadequately characterized.

We can apply our approach to track changes in protein phospho-signals throughout the cell over time. These changes typically correlated with cellular events observed via IF. For example, we identified context-specific ciliary protein phosphorylations that are normally either undetectable or invariant using phospho-proteomics of asynchronous cells. We identified phosphorylations of known regulators of ciliary vesicle traffic that spike right when RAB34 recruits to the basal body (∼7-8hr, and again at ∼18-20hr): RAB34 [S241, S244] and MYO5A [S1652]. These and many other instances show the temporal consistency for when modifications to these proteins are occurring and when these proteins are known to function, both in the literature and based on our microscopy timeline (Fig.2A).

Having identified strong correlations between the timed appearance of specific phosphoproteins and known cellular events linked to these proteins as observed by IF (Fig.2A), we applied this framework to generate hypotheses about the dynamics of the general cell program during the G_0_ transition. We performed correlation analysis of all the temporal phospho-signatures to identify those with similar patterns. We then binned them to identify which cellular compartments/functions are undergoing changes between different time windows. In brief, data from 3 biological replicates were analyzed to identify trends in their temporal phospho-signatures. Phosphopeptides were sorted into 30 different groups and plotted in a heat map (Fig.5A). Each individual row of the map corresponds to a single phospho-signature and its change in relative abundance over time, with the x-axis indicating the time and consistent stage of ciliogenesis (based on IF). Levels of phosphopeptide abundance are represented as a gradient of blue to yellow, with yellow being the highest abundance. The left of the map color codes the clusters in which we binned phosphosites that share the same temporal signatures of fluctuations in abundance (Fig.5A).

**Fig.5.**
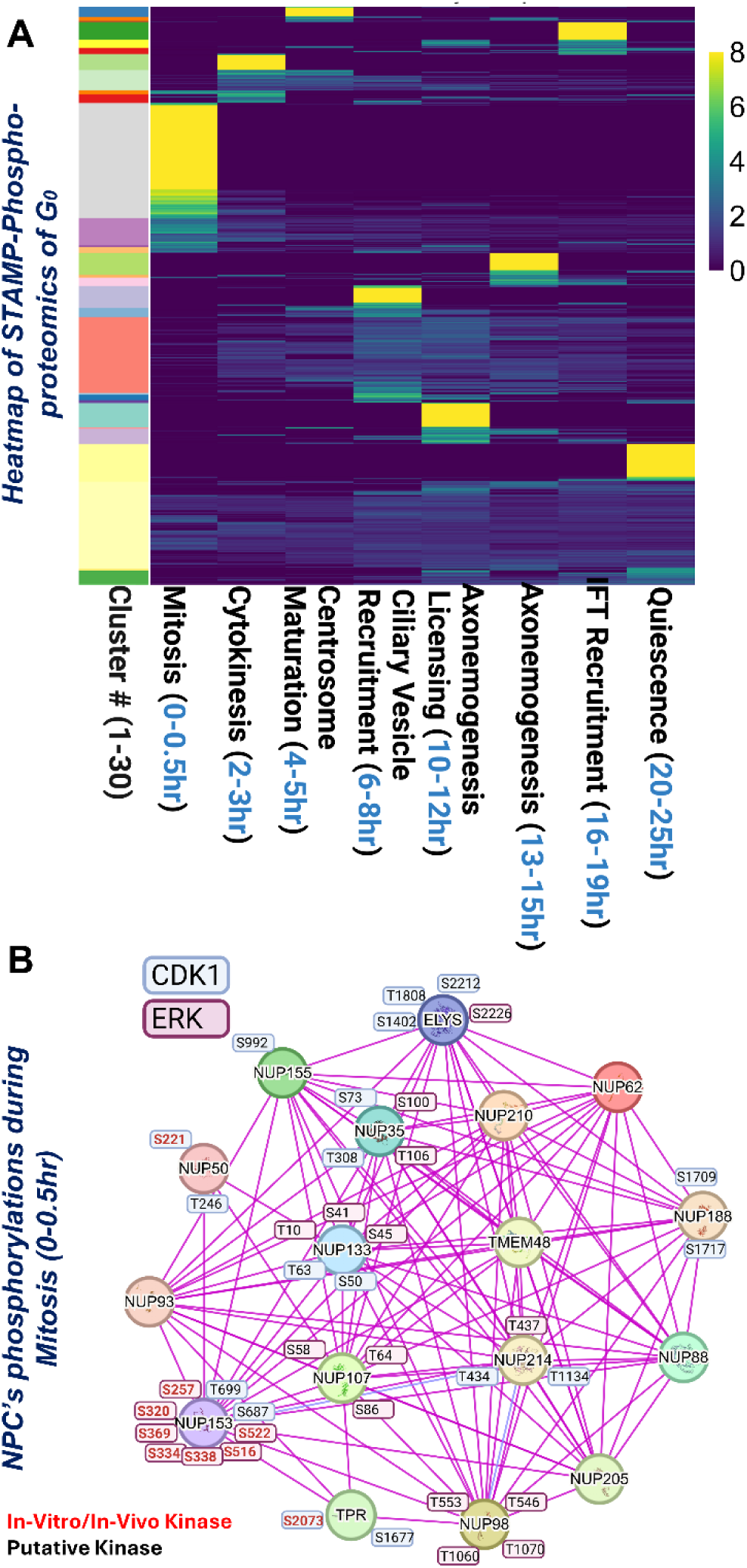
Phospho-proteomics reveals key signatures reflective of dynamic changes in the cellular landscape during the transition into quiescence. (A) Clustered heat map of the normalized intensity of phosphopeptides after STAMP, as a compilation of 7 separate experiments with multiple technical replicates per each experiment. The color gradient goes from low (dark blue) to high (yellow) as indications of fold change, as shown on the right. On the bottom left, colored boxes indicate individual clusters in which phosphopeptides were binned based on similarities in the pattern of the intensity/abundance of individual phosphopeptides at different time points after seeding, which we term their “phosphosignatures.” This heatmap was generated using R programming software, performing correlation analysis to cluster phosphopeptides with similar phospho-signature patterns. The left shows the identifier of each cluster, which can be used to go to the Excel sheets and identify which phosphopeptides are within your cluster of interest. The bottom axis shows the time window and the ciliogenesis stage that occurs in this window. (B) Schematic representing NPC complex and phosphorylation events happening during mitosis. The phosphorylation boxes are color-coded as purple boxes for ERK substrates and blue boxes for CDK1 substrates. Black text means the indicated kinase is putative, and red indicates that the indicated kinase has been tested and verified based on the Phosphosite.(*1*) *Created in BioRender. Turn, R.* (*2025*) https://BioRender.com/o71n224.

In this way, we can look at the data from multiple angles and determine whether: 1) finding a protein of interest, identifying the cluster(s) its phosphopeptides are in, and identifying putative functions; or 2) analyzing any individual cluster of interest to gauge what cell functions are acting in concert in each time frame during G_0_ transition. In the latter case, we performed gene ontology (EnrichR) (Fig.S5) and Cytoscape analyses of individual clusters to identify whether proteins that are undergoing phosphorylations in specific windows share known cellular functions (Fig.S5). These data were used to generate a temporal-spatial map of the phospho-signatures emerging through G_0_ progression and linked to specific cellular locations or processes.

Many of the functions identified via gene ontology occurred within the predicted time frame, based on our temporal-spatial mapping via IF. For example, we observed major changes in phosphorylations of mitotic machinery in 0-0.5hr and endosome traffic proteins at 8hr and 12-20hr. We also identified trends that pointed to key functional windows, based on a common “time-stamp” for phosphorylation of a linked set of factors. For example, we noted a class of phosphorylations that only emerged after cilia had fully formed (*i.e*, were at full length and presented with ciliary GPCRs), increasing at ∼20hr and remaining elevated. These include CEP170[S1198], CP110[S372], and CEP97[S114]. Note that CP110 functions early-on to prevent spurious ciliogenesis, and indeed CEP97 is the inhibitor target of CP110, and CEP170 is critical to traffic regulators to the centriole. Yet, no function has been reported for these factors later in the G_0_ program. One possibility could be that the CP110 inhibitor program is only established late in G_0_ progression, once ciliary signaling is established, to limit further ciliary assembly. Together, the combination of IF, phospho-proteomics, and known function mapping enables the generation of innovative, testable hypotheses as to the dynamic landscape of metaphase through G_0_ transition. In the sections below, we provide examples of how we tested some of these hypotheses generated from phospho-proteomics and IF.

### Regulation of NPC machinery and autophagy machinery as revealed by STAMP phospho-proteomics

We also tracked phosphorylations for known mitotic regulators to see if the time-stamped phospho-signatures of proteins that we detected correspond with their known functions and, therefore, predicted time-stamps. In Fig.5B, we provide a STRING PPI plot of all the interacting components of the NPC complex that were phosphorylated based on our phospho-proteomics analysis. We specifically traced putative and validated ERK and CDK1 substrates on the nuclear pore complex (NPC), which have known functions in NPC disassembly during mitosis (*44–48*). ERK and CDK1 are key kinases that hyper-phosphorylate specific nucleoporins such as NUP35, NUP153, NUP188, and ELYS(*44*). All the phosphorylations annotated in the plot occur specifically within the 0-0.5hr window, consistent with their mitotic function. Furthermore, all the phosphorylations shown here are either predicted (black font) or known (red font) to be CDK1 or ERK substrates (Fig.5B). Future work is needed to determine whether ERK or CDK1 are the true upstream kinases for the putative sites, though the common motifs, the high scoring based on the Cantley screen, and the timing of when they achieve peak abundance all support this hypothesis. Note that some of the NUPs shown in this STRING plot are unphosphorylated in this time window. Instead, these NUPs typically either were phosphorylated in cytokinesis, or their phospho-signatures remained constant throughout the time course. Beyond NPC machinery, we also detected phosphorylations of Golgi fragmentation regulators in mitosis (*e.g*, GOLGA1[S767], GM130[S37], BET1[S48]), consistent with Golgi fragmentation and later reassembly during the G_0_ transition. This coincides with our IF data (Fig.1D).

Note that known phosphorylations that mark other states of cell cycle were undetectable under these conditions, including but not limited to G_1_ (RB1[S780/S795] and RB1[S807/S811]), G_1_/S transition (RB1[S612/S672] and p27[T187]), S phase (H3[S10]), and G_2_ (Cdc25[S216]). These signals are normally very pervasive and easy to detect in standard phospho-proteomics of heterogeneously cycling cells, and even in asynchronous ciliogenesis samples. This suggests that STAMP phospho-proteomics specifically tracked the transition of cells from mitosis to quiescence, consistent with the IF data in Fig.1. Further, these observations also support our empirical findings that the 20-minute recovery in serum-enriched medium after nocodazole treatment did not alter cell cycle progression or quiescence. Reports in the literature describe specific phosphoproteomic signatures that emerge following nocodazole toxicity or exposure(*28*). However, these nocodazole-associated signatures are absent in our data. Interestingly, several validated phosphorylations, such as STAT3[S727](*49*), PLK1[S210](*47, 48*), PX2[S121](*50*), and JUN[S63](*51*), have been reported as potential long-term signatures that emerge in cells after nocodazole treatment. After performing the STAMP protocol in RPE cells, we observe that these phosphorylations are present at low levels during metaphase, increase significantly during cytokinesis, and then decline after cell division, remaining at low levels (Fig.S6). These findings suggest that these signatures may represent cytokinetic signals, as their peak expression coincides with cytokinesis rather than nocodazole exposure. Perhaps cells that sustain nocodazole toxicity retain features of the cytokinetic program.

STAMP phospho-proteomics was especially useful for temporal-spatial mapping of transient time-stamped phospho-signatures for regulators of established signaling pathways. For example, we mapped temporally-restricted phosphorylations to key components of the mTOR autophagy and protein synthesis pathways and observed key hierarchical signaling. In Fig.6, we mapped phosphorylations that we annotated via STAMP phospho-proteomics onto the known regulators of autophagosome maturation, noting the time frames in which these phosphorylations peaked in abundance. The color of the phosphorylation shown indicates either the known (based on the literature) or putative kinases (as predicted based on human kinome substrate specificity algorithm)(*52*) that mediate these functions. The temporal mapping of phosphorylations of known function in the mTOR pathway proved especially valuable for identifying context-specific, timed, and site-specific signaling in distinct cellular locations.

**Fig.6.**
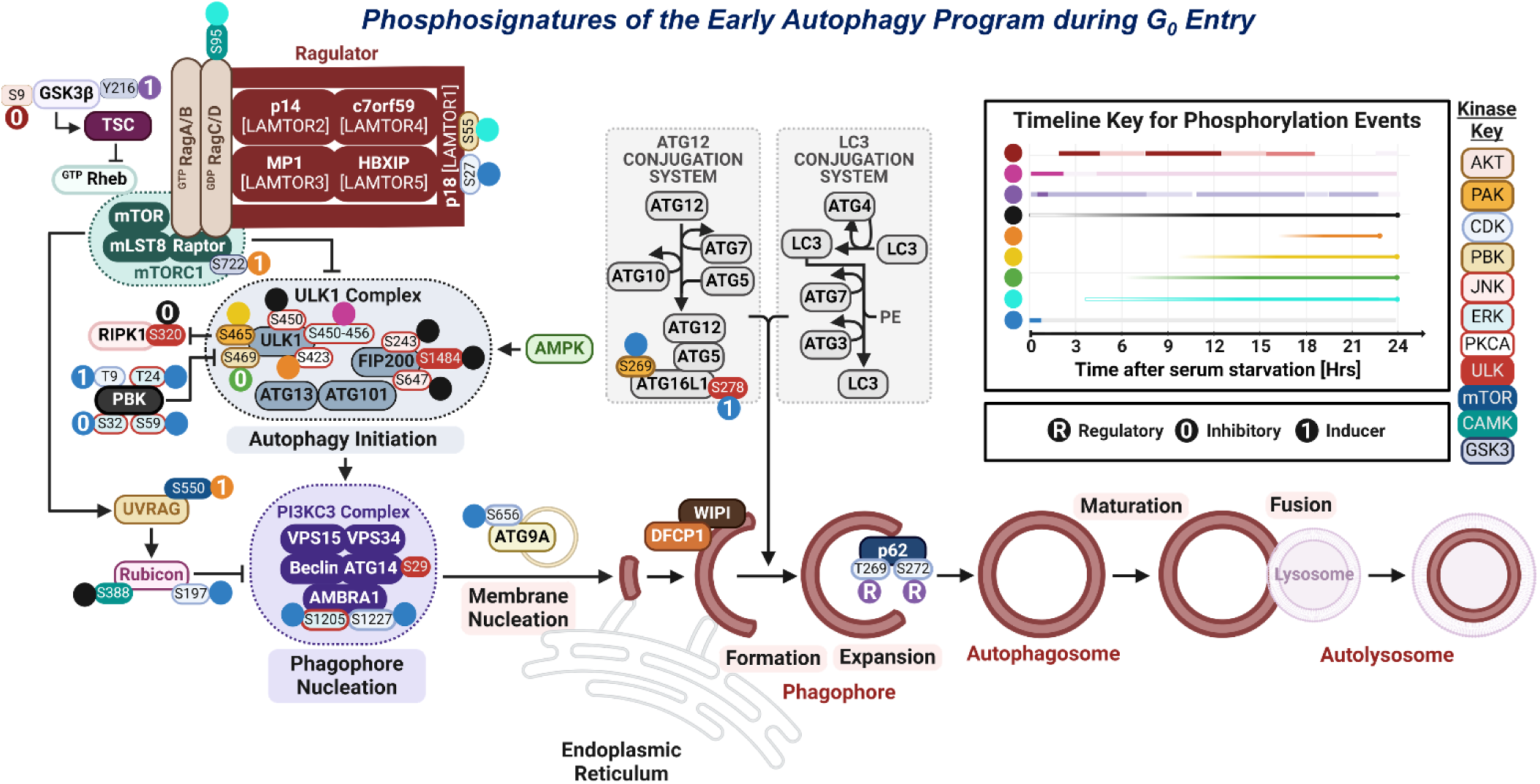
Mapping the autophagy program during G_0_ entry using STAMP. Key examples of the vast array of temporally regulated phosphorylations of known autophagosome maturation machinery, as identified via STAMP. Each phosphorylation is annotated with a colored dot indicating its pattern of abundance over time (see chart on the upper right), and the color of the box surrounding the phosphorylation indicates the putative/known regulatory kinase (see Kinase Key on the right of the dot timeline). Created in BioRender. Turn, R. (2024) BioRender.com/b48o404.

As one example, GSK3β is a negative regulator of mTORC1, promoting conditions that favor autophagy, especially during energy deprivation or stress. GSK-3β activation is markedly elevated during cytokinesis, as evidenced by high levels of the [Y216] activating phosphorylation site and negligible levels of the [S9] inhibitory phosphorylation. This pattern suggests a state of mTORC1 inhibition and autophagy induction, alongside GSK-3β’s crucial role in regulating microtubule dynamics during this process. Following cytokinesis, GSK-3β activity decreases; however, during ciliogenesis, we observe a further decrease in GSK-3β[Y216] activating phosphorylation at two additional time windows: at ∼9 and later ∼18 hrs post-serum starvation, coinciding with increased inhibitory [S9] phosphorylation. The interplay of these phospho-signatures indicates a shift in regulatory mechanisms as cells progress into quiescence, suggesting two waves of autophagy during the transition (Fig.6).

To further explore the potential function of autophagy in multiple stages of G_0_ progression, we performed immunofluorescence of synchronized cells and tracked lysosome distribution over time (using LAMP1 as a classic marker of lysosomes) (Fig.7). We observed that, over time, LAMP1-positive puncta diminish and that the remaining pool of lysosomes clusters around the basal body at 6hr. This is consistent with the time frame in which GSK3-β [Y216] activation signal returns. The pool of lysosomes remains at the basal body even after the primary cilium has formed, though we do observe that the lysosome pool after ciliogenesis is less compact (Fig.7). The dynamic clustering of lysosomes around MTOCs has been described(*2*), but no previous studies link this to a specific stage in G_0_. This temporal ordering of phosphorylations allows us to resolve time and context-specific kinase activities and functions that would have been impossible to detect without STAMP. Exploration of signaling pathways and their dynamic functions throughout G_0_ transition and in response to stimuli would be a rich field for future investigation.

**Fig.7.**
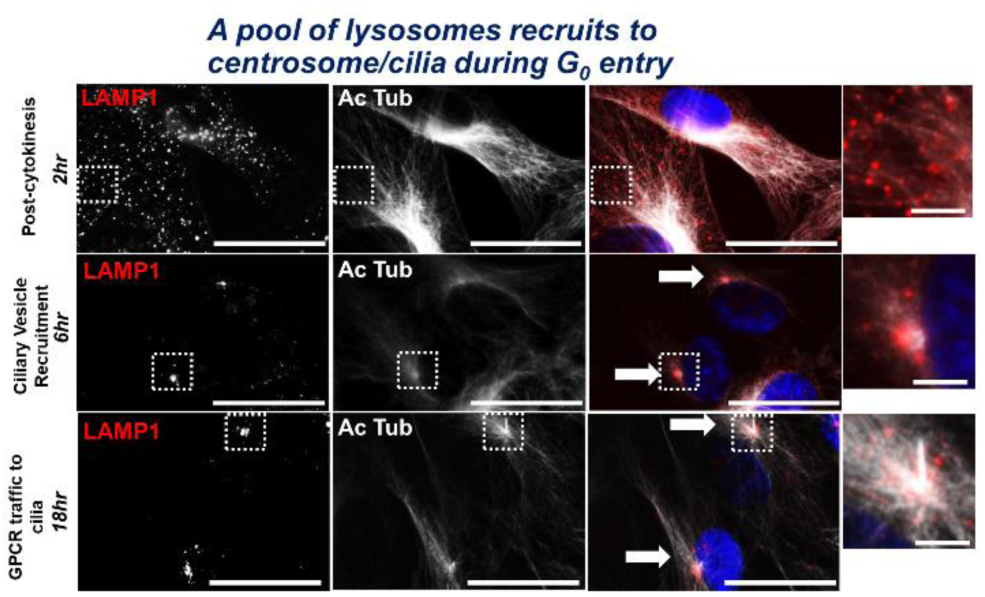
Representative images of LAMP1-positive lysosome distribution throughout the G_0_ program. Images were collected via immunofluorescence of G_0_ synchronized cells via widefield microscopy at 60x magnification, full scale image error bar= 20μm, insert=1μm. Cells were stained for LAMP1 (red), acetylated tubulin (white), and DAPI (blue).

### Defining detailed, time-stamped events driving ciliogenesis via live cell imaging

While many cellular events can be monitored by simple static microscopy images, a number of dynamic cellular events are best visualized by live cell imaging, which allows us to determine the time-stamp on more dynamic changes in cellular morphology and protein localization. As proof of principle, we performed STAMP synchronization using RPE cells stably expressing RABL2-GFP to mark distal appendages and IFT88-mCherry to mark IFT-B trains. We co-stained with SiR-Tubulin to track axonemogenesis. We then performed multi-field, z-stack confocal imaging at 40x magnification, every 8min over a 24hr time course.

Though our initial focus was on ciliogenesis as a marker of the transition to quiescence, live cell imaging also provided insight into other aspects of cellular function. Within minutes after seeding, cells proceeded to cytokinesis, which finished within 2-3 hrs (Fig.8A), as made evident by the tubulin signal. Interestingly, at ∼2hrs, a short-lived phenomenon occurred in which tubulin accumulated in a large “barrel-like” structure surrounding the centrosome (2:33 time point in Fig. 8A). These tubulin-rich barrels were encircled by IFT88 vesicles (Fig.8A, Fig.S7). Less than an hour after forming, this tubulin structure dissipates, and the IFT88-positive vesicles become evenly distributed throughout the cell (∼4.5hr). Later (12hr), IFT88 positive vesicles aggregate around the centrosome (Fig.8B), with no sign of the return of the tubulin barrel intermediate. IFT88 later extends up the length of the growing cilium, consistent with IFT-B complex entry. Thus, the high degree of cell synchronization revealed a key, currently unexplained, transient phenomenon of tubulin and IFT88 aggregation at centrosomes and allowed temporal resolution of key events.

**Fig.8.**
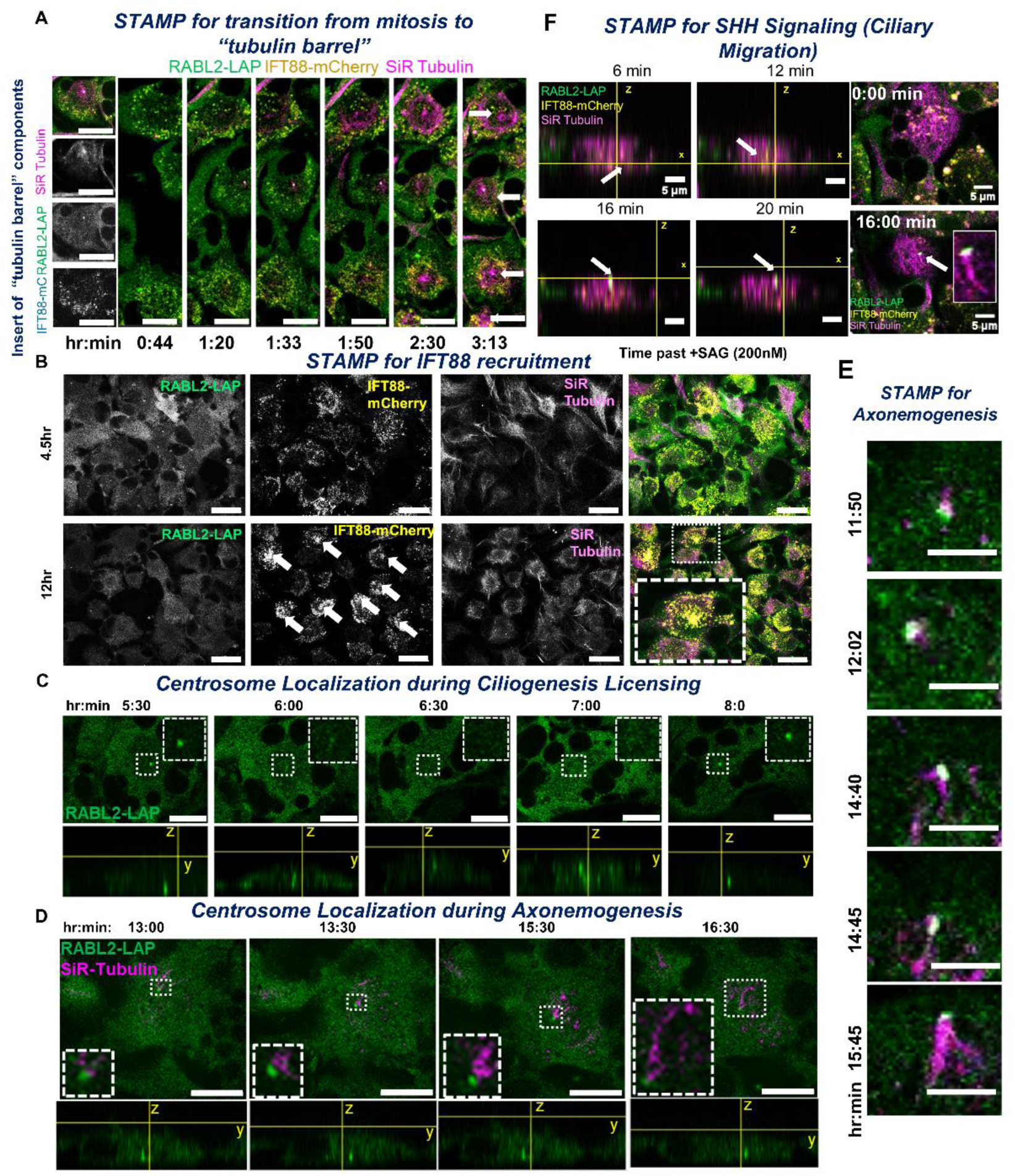
Live cell imaging reveals transient intermediate states in cell morphology and signaling during transition into quiescence. Live cell time lapse imaging of STAMP synchronized, fluorescently tagged (IFT88-mCherry, RABL2-LAP(GFP)) and SiR-tubulin stained RPE cells over a 24hr-long time course. Images were collected at 40x magnification with z-stacks every ∼2-8min, with multiple fields per experiment. These experiments were performed in multiple replicates, with representative images included. Key examples of events captured via time-lapse imaging include the following: (A) a post-mitotic tubulin barrel at ∼2hr, Scale = 10μm (B) IFT88-positive vesicle recruitment to the centrosome before entry into the cilium at ∼12hr, Scale = 20μm (C) dynamic basal body localization throughout ciliogenesis licensing, Scale = 10μm, and (D) static basal body localization during axonemogenesis. Scale = 10μm (E) axonemogenesis within a ∼1hr time frame; Scale = 1μm and (F) ciliary migration within minutes of SHH induction. Scale = 5μm.

We also tracked the dynamic localization of the centrosome during different stages of the G_0_ program (Fig.8C)(*53*). After cells adhered to the plate, centrosomes localized basolaterally for the first ∼6hr. The basal body then translocated to the apical surface between 6-8hrs, at which point it sunk basolaterally again. This window coincides with the ciliary vesicle recruitment and early ciliogenesis licensing.

On the other hand, basal body localization appeared to be more static later in the ciliogenesis program, specifically during axonemogenesis (Fig.8C-E). Essentially all cells began ciliogenesis within a 2hr window, starting at ∼11hrs (Fig.8C). Live cell imaging of synchronized cells also allowed us to determine the duration of specific cellular events on a cell-to-cell basis. For example, even though axonemogenesis in any given cell takes ∼90min to complete,(*22*) axonemogenesis across the cell population occurred within a 4hr window (13-17hr; Fig.8D-E). While tracking axoneme extension via live cell imaging with SiR tubulin as a marker of axonemes, we observed that the basal body remained relatively in the same focal plane, sitting adjacent to the nucleus (Fig.8D). It would be interesting to repeat these studies using membrane markers to observe whether these cilia are internal or whether we can detect extension of the axoneme outside of the cell.

We also posited that live cell imaging with STAMP provides a valuable approach for studying ciliary signaling. We stimulated the Sonic Hedgehog pathway with SAG (a synthetic Smoothened agonist, 200nM), and within 8min of stimulation we observed basal to apical translocation of primary cilia and increased accumulation of IFT88-mCherry particles at the basal body (Fig.8F). Thus, we can identify key changes in the cell related to cilia and ciliary function and employ MS and fixed cell imaging at specific time points to pursue mechanistic insights.

### Applying PIT-STAMP to dissect temporal-spatial mechanisms

We predicted that cell synchronization would decrease background “noise” and would allow us to dissect the effects of individual regulators on different features of G_0_ progression. To do so, we developed an approach called <u>P</u>ulsed Inhibitor <u>T</u>reatment (PIT)-STAMP, which involves the treatment of synchronized RPE cells at different times to determine the acute effects of loss of key regulators on the G_0_ program. We began by using PIT-STAMP to test the hypothesis that translational control helps establish the G_0_ program and ciliogenesis. Phospho-proteomics and Gene Ontology revealed phosphorylations of transcription and translation machinery at different stages of G_0_ transition (Fig.S5). Examples of such machinery include RSK2[S369], PRAS40[T246], p70S6K[S447], NCL[S67], GAPDH[T177], YTHDF1[T202, S217], FXR2[S601, S603], SSB[S92, S225], and LARP4B[S736, T732]-all of which share the time-stamp of peak intensity at 4hr. The activating phosphorylation RSK2[S369], in particular, is known for driving its function in transcriptional and translational control downstream of ERK(*54*). When we looked again at mTOR pathway phosphorylations, key modifications peak at 4hr, including known regulators of transcriptional and translational control.

We tested our hypothesis by adding protein synthesis inhibitor cycloheximide (10μg/mL) for 1hr-long intervals starting at 4hr, 6hr, 10hr, and 16hr post-seeding, followed by washout and return to serum free medium to allow collection of later time points. While ciliation was unchanged in cells treated with cycloheximide at the later time points, ciliogenesis was completely ablated in response to cycloheximide treatment at 4hr, even though the drug was removed at 5hr. This suggests that protein synthesis early in G_0_ transition is essential for ciliogenesis. The role(s) of later windows of translational control in establishing the G_0_ program is unknown but would be an exciting direction to study. Altogether, these data establish a paradigm for future research into which proteins are synthesized in which G_0_ transition windows and their respective functions.

PIT-STAMP also can be used to assess effects of inhibitors on regulators of specific pathways. We again use ciliogenesis as an example. We dosed cells with dynarrestin, a cytoplasmic dynein inhibitor that prevents microtubule association and, therefore, blocks retrograde ciliary transport(*55*). We added dynarrestin (25μM) at the same 1hr intervals described above for cycloheximide, followed by washout and fixation at 24hr (Fig.9A-C). Treatment with dynarrestin only inhibited ciliogenesis (from ∼80% to ∼5% ciliation) at the latest (16-17hr) window, or during axonemogenesis but before GPCR recruitment. A partial (∼40% ciliation), though not significant, reduction of ciliogenesis occurred when added in the 10-11hr window. This is consistent with cytoplasmic dynein and IFT being required to extend the axoneme and traffic key cargoes into growing cilia (Fig.9B-C).

**Fig.9.**
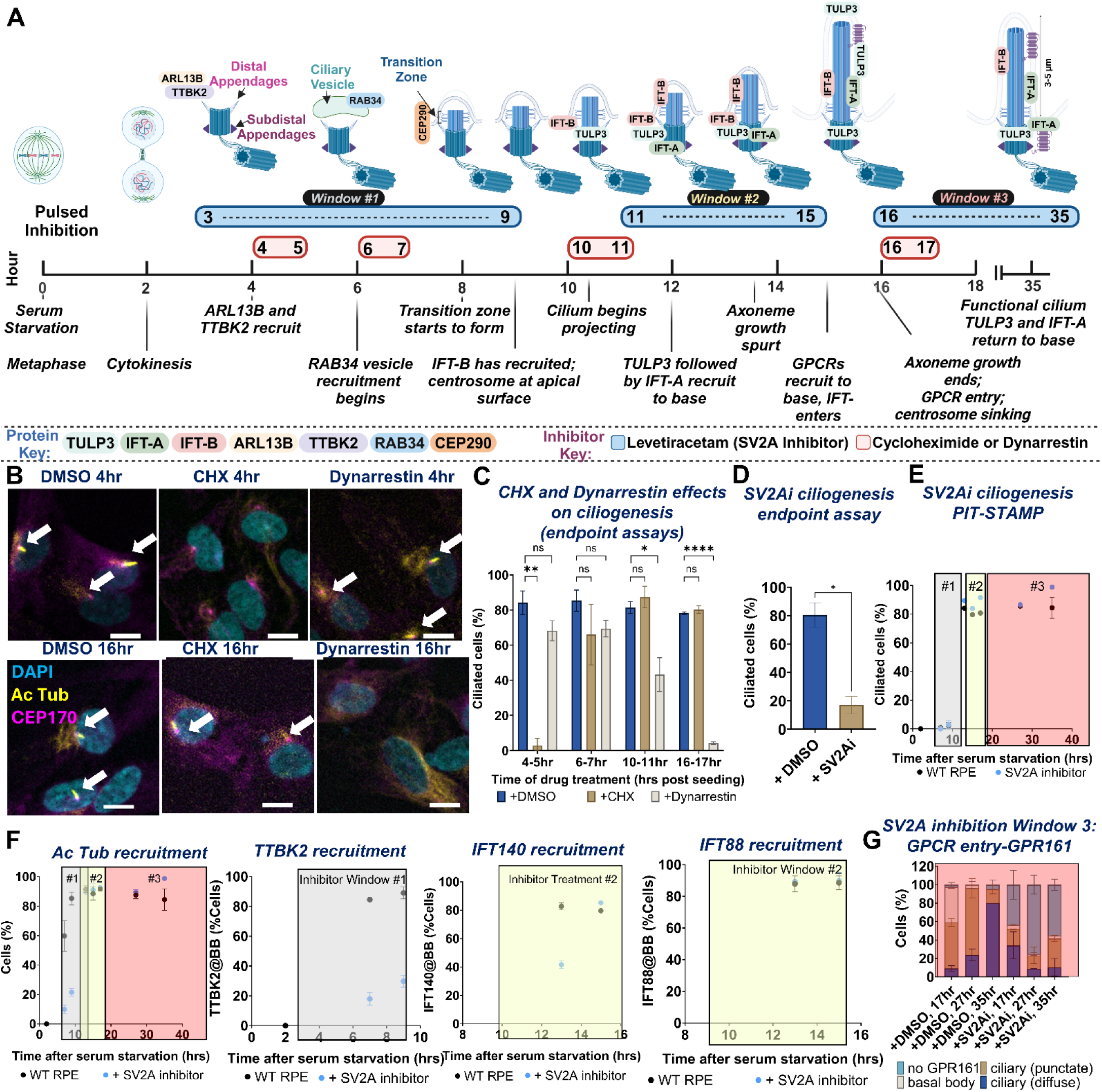
Mapping the regulatory kinases and phosphatases that drive the intracellular landscape of G_0_ transition. (A) Timeline mapping when individual drug treatments were performed for PIT-STAMP, as mapped onto ciliogenesis as described previously in Figure 2. Red windows indicate when cycloheximide and dynarrestin drug treatments were performed (1hr spike-ins before washout, followed by endpoint fixation at 24hr), while blue windows refer to time frames in which SV2A inhibitor (levetiracetam) was added (long-term drug treatments but collecting intermediate time points in between each window to track intermediate steps). Created in BioRender. Turn, R. (2025) https://BioRender.com/x79z074 (B) Representative images showing the different effects of cycloheximide (CHX) and dynarrestin treatments at various PIT-STAMP’s windows (4hr vs 16hr window treatment) compared to DMSO control. Images were collected via widefield at 20x magnification using ImageXpress software. Blue= DAPI, magenta = CEP170, yellow = acetylated tubulin. Scale bar = 5μm. C) Quantification of ciliation rate in dynarrestin and CHX PIT-STAMP treated cells versus DMSO control. (D) Quantification of % ciliation in WT RPE with and without SV2Ai (levetiracetam) treatment at 4hr post-seeding of synchronized cells, without washout. (E-G) Quantification of IF images of PIT-STAMP for SV2A inhibitor treatments on ciliogenesis and ciliary function. Different markers were quantified for different treatment windows: Window 1-TTBK2, mitotic indices; Window 2-IFT140, IFT88, ciliation; Window 3-GPR161, ciliation. Experiments were performed in replicate, N=100 cells per replicate. Grey box= Window 1, Yellow Box = Window 2, Red box= Window 3. All experiments were performed in replicate, counting N=100 cells for each phenotype after IF. Error bars = SEM. Statistics performed via One-Way ANOVA, with *= 0.05, **=0.01, ***=0.001, ****=0.00001. Plots were generated with GraphPad Prism Software.

### Use of genetic modifications to aid in the ordering and dissection of specific processes

We predicted that STAMP can also be used to dissect mechanisms of known but not fully characterized regulators of cell functions through performing the same procedures but in the context of genetic perturbation (*e.g*, CRISPR KO). To test this, we used STAMP to systematically phenotype ciliogenesis defects in ARL13B, TTBK2, and IFT52 KOs versus WT. Though each of these proteins are clearly important in ciliary function, the temporal-spatial mechanisms by which these factors drive ciliogenesis have yet to be elucidated.

Synchronization and IF of these three ciliogenesis defective cell lines revealed key differences concerning the nature of their ciliogenesis-defective lesions. Each of the three KO lines failed to ciliate (WT 80-90% versus ∼0%; Fig.10A). However, the three KO lines diverged from each other in acetylated tubulin recruitment to the basal body and RAB34-positive ciliary vesicle recruitment. RAB34 recruitment in WT RPE occurs in two windows: peaking at ∼7.5hr and again at ∼20hr. TTBK2 KO cells recruited a first wave of RAB34 positive vesicles to the basal body (Fig.10A). However, the second wave of RAB34 recruitment and GPR161 recruitment is lost in TTBK2 KOs. IFT52 KO showed both a reduced and delayed first wave of RAB34 recruitment, peaking at only ∼40% of basal bodies positive for RAB34 (10hr), suggesting an earlier function in licensing (Fig.10A). As observed in the TTBK2 KOs, IFT52 KOs do not have a second wave of RAB34 recruitment but still have comparable acetylated tubulin recruitment to the basal body with WT. IFT52 KO cells show reduced GPR161 recruitment to basal bodies and defective intraflagellar traffic. In ARL13B KO cells, the first wave of RAB34 recruitment to the basal body is similar to WT, peaking at ∼80% of cells at ∼7.5hr, but there was no second wave. Instead, we observed a striking accumulation of RAB34 vesicles into what appears to be “aggregates’’ throughout the cell that began at 4hr and continued to enlarge throughout the rest of the time course (Fig.10B). ARL13B KOs also had defective acetylated tubulin recruitment, presenting with normal formation of the acetylated tubulin puncta at 7.5hr which later dissipated at 10hr, with only ∼20% of basal bodies being positive for acetylated tubulin by the end of the time course. Thus, the initial wave of ciliary vesicle recruitment is independent of TTBK2 but dependent/interconnected with ARL13B and IFT52/IFT-B function. Also, these data lead one to question whether RAB34-positive vesicle recruitment licenses acetylated tubulin recruitment and eventual extension of the axoneme. The degree to which each of these proteins is directly or indirectly involved in the function of these markers is currently unclear, but STAMP provides an optimized approach for future work pursuing mechanistic details. Altogether, STAMP combined with genetic manipulations serves as a powerful tool for dissecting the underlying mechanisms by which proteins regulate cellular processes and provides testable hypotheses for future studies.

**Fig.10:**
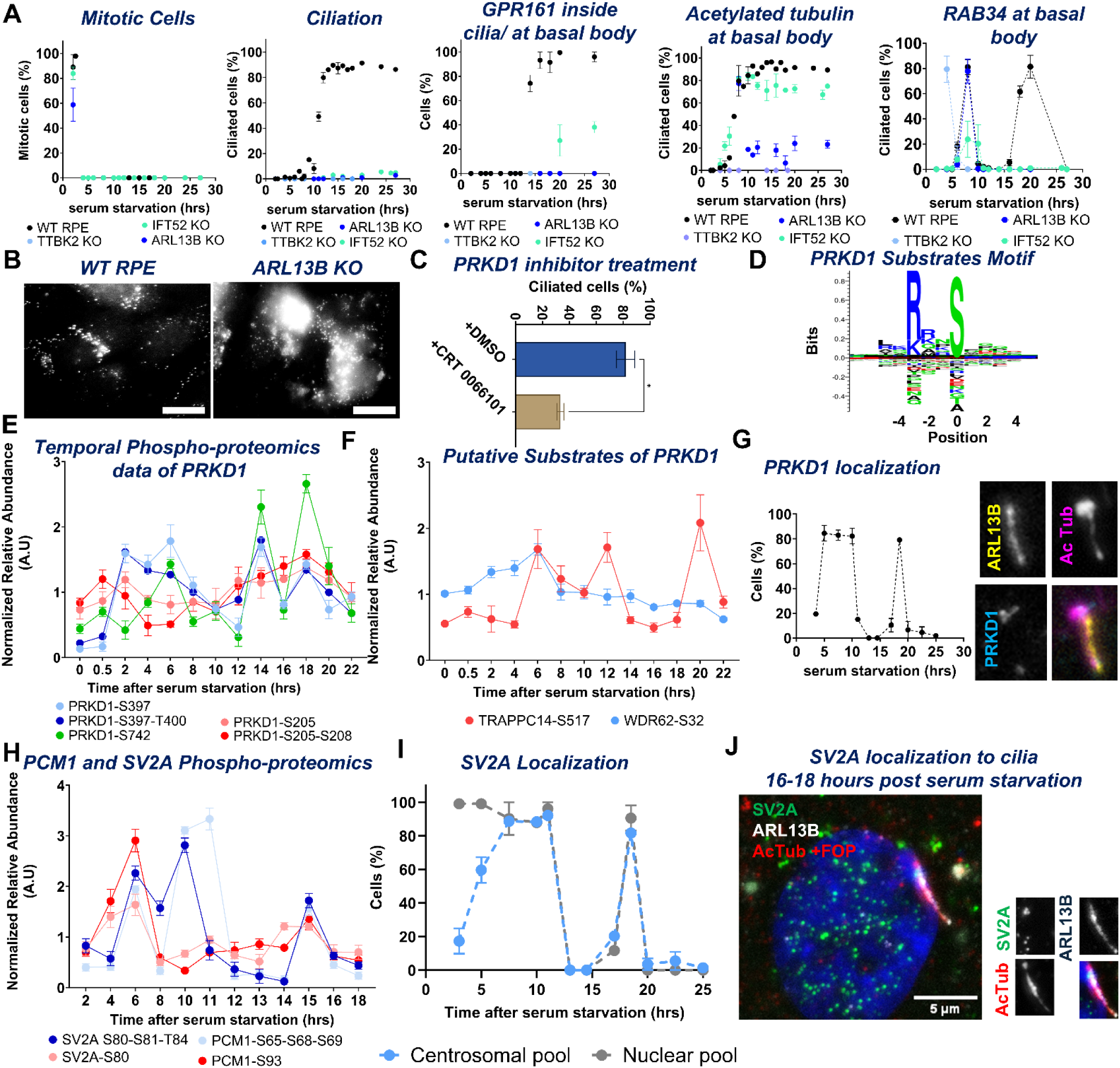
Applications of STAMP to decipher key functions and regulators of G_0_ transition. (A) Quantification of IF markers to compare rates of ciliogenesis and G_0_ entry between different KOs: ARL13B vs TTBK2 vs IFT52 KO. Experiments were performed in replicate. (B) Representative images of Rab34 vesicle morphology in WT versus ARL13B KO RPEs 24hr post-initiation of STAMP. Z-stacks were collected via widefield microscopy at 60x magnification. Scale = 20μm (C) Quantification of PRKD1 inhibition effect on the ciliogenesis, (D) Motif of PRKD1 substrates from STAMP-Phospho-proteomics for sites with higher score than 99 percent of all putative phosphorylation sites in the phosphoproteome based on Cantley’s algorithm. (E) Normalized abundance fluctuation of phosphorylations on PRKD1, and (F) two putative PRKD1 kinase substrates including TRAPPC14 [S517] and WDR62 [S32], (G) Representative images, quantification of localization of PRKD1 to cilia/centrosome. Images were collected via widefield microscopy at 60x magnification. (H) N-terminus phosphorylations on the SV2A shows similar patterns of phosphorylations on known ciliary proteins PCM-1, (I) Quantification of localization to cilia/centrosome, for SV2A, as another regulator of ciliogenesis, and (J) it’s representative images. Images were collected via widefield microscopy at 60x magnification, error bar= 5μm. Quantification was performed in replicate. Note that SV2A inhibitor was used above in Fig.9 to determine whether it also played a role in ciliation. Phospho-signature mapping was performed to measure normalized intensity of different phosphopeptides for multiple replicates. For IF quantifications N = 100 cells per replicate.

### Using STAMP to identify key regulators of cellular functions in G_0_

Based on the phospho-proteomics and correlation clustering of time-stamps that we performed above (Fig.3A), we hypothesized that critical regulators functioning in specific stages of ciliogenesis would share similar phospho-signature fluctuation patterns and time-stamps with known regulators of that function. We already identified known regulators of ciliogenesis and G_0_ that exhibited phospho-signature patterns that correlated with the timing of their known functions and localization via IF, based on our ciliogenesis timeline (Fig.2B). Of the ∼15,000 phosphorylations detected during the time course, ∼7500 phosphosites showed dramatic changes in intensity that we later binned based on correlation analysis of phospho-signature patterning. Here we discovered two examples of key ciliogenesis regulators identified from these analyses. Protein Kinase D1 (PRKD1) (Fig.10C) and Synaptic Vesicle Glycoprotein 2A (SV2A) (Fig.10D). PRKD1, which regulates TGN signaling(*56*), revealed transient phospho-signatures that suggested activity during ciliogenesis/G_0._ In this study, we demonstrate how STAMP can be utilized to generate mechanistic hypotheses for cellular events in a systematic manner. To explore whether the loss of PRKD kinase activity alters ciliogenesis, we used PIT-STAMP on wild-type RPE cells and treated them with the general PRKD inhibitor CRT 0066101 (5μM) starting at 4 hours post-cytokinesis, maintaining the treatment throughout a 24-hour time course (Fig.10). Under these conditions, ciliogenesis was reduced from approximately ∼80% to ∼35%. However, due to the non-selective nature of kinase inhibitors, it is challenging to pinpoint which PRKD isoform is involved in ciliogenesis. Complementary data from STAMP phospho-proteomics, however, suggests that PRKD1 is the primary isoform involved. We found no unique phosphorylation sites for PRKD3, while phosphorylation events for PRKD2 at sites [S197, S198] were more abundant during mitosis and cytokinesis, suggesting its role in these processes (Fig.S8). Several time-resolved phospho-signatures for PRKD1 were identified, including the PRKD1 autophosphorylation stimulatory site [S742] and doubly phosphorylated [S397, T400] and [S397] sites, all of which increased during vesicle recruitment and GPCR trafficking at the 6- and 18-hr time points (Fig.10C). Additionally, PRKD1 localized to the basal body in two distinct waves, during the 5-10hr and 17-20hr windows, consistent with its role in vesicle recruitment and IFT. This timing of phosphorylation events correlates with changes in PRKD1 localization. We also identified putative PRKD1 phosphorylation sites based on Cantley analysis, such as the transient phosphorylation of TRAPPC14 [S517] during vesicle recruitment, correlating with PRKD1 phosphorylation events. TRAPPC14, a component of the TRAPPII complex, regulates Rabin8 preciliary trafficking and ciliogenesis(*57*). Furthermore, phosphorylation of WDR62 [S32] exhibited similar intensity trends, with favorable sequences for PRKD1-mediated phosphorylation. Previous studies revealed that WDR62 point mutations perturb recruitment of CENPJ to the basal body and caused loss of axonemal IFT88, which is required for ciliary protein trafficking(*58*). While these findings require further validation to exclude potential involvement of other kinases and to confirm the functional roles of the phosphorylation sites, these STAMP data effectively highlight its capacity to generate testable hypotheses.

We provide yet another proof of principle for STAMP as a tool for identifying dynamic regulators in the case of SV2A, an 83kDa transmembrane protein and galactose transporter that regulates neurotransmitter release and is a major target of anti-epileptic drugs.(*59–61*) The temporal phospho-signatures of SV2A were very similar to those of PCM-1, a regulator of ciliogenesis that coordinates centriolar satellite remodeling to allow ciliation and is also critical for ciliary vesicle docking.(*62–65*) Specifically, SV2A is triply phosphorylated on [S80, S81,T84], resembling PCM-1 phosphopeptides, including PCM-1[S65, S68, S69] and PCM-1[S1765, S1768, S1776] (Fig.10D). These triple phosphorylations contain similar motifs suggestive of potential activation by the same kinase, likely through priming kinase activity (or a hierarchy of phosphorylations in which the first phosphorylation serves as a recognition motif for following phosphorylations). Based on these correlations, we predict that SV2A plays a role in vesicle recruitment to cilia. To test this hypothesis, we tracked SV2A distribution temporally in synchronized RPE cells (Fig.10D). We observed transient localization of SV2A to the basal body between 5-11hr and at what appears to be the transition zone of primary cilia at 17-18.5hr (Fig.10D). These time windows correspond to the window where we see RAB34-positive vesicle recruitment, based on our temporal-spatial map (Fig.2). If SV2A is playing a direct role in ciliary vesicle traffic to drive ciliogenesis, we predicted that inhibition of SV2A would inhibit ciliogenesis specifically at stages associated with vesicle traffic. To test if SV2A plays a role in ciliogenesis, we inhibited SV2A using the highly selective SV2A inhibitor/ antiepileptic drug levetiracetam (50μg/mL) by treating synchronized cells at 4hr and maintaining the drug treatment throughout the imaging window (Fig.9D). We observed that ciliogenesis decreases from ∼80% to ∼20% upon SV2A inhibition. Furthermore, the few cilia that did form in SV2A-inhibited cells had defective GPR161 ciliary entry. We then blocked SV2A activity using the highly selective SV2A inhibitor/ antiepileptic drug levetiracetam (50μg/mL) during 3 different time windows without washout (Fig.9A,E) to see if we could pinpoint the lesion. Levetiracetam did lead to reduced TTBK2 recruitment and localization of acetylated tubulin after treatment in the 5hr window Treatment between 11-15hr did not delay IFT-B recruitment, but it did delay IFT-A recruitment. Also, SV2A inhibition from 16-35hr delayed GPR161 entry into primary cilia, consistent with roles for SV2A in coordinating ciliary traffic (Fig.9E-G). Ciliation rates were unaffected by SV2A inhibition in Window #2 and Window #3. These data together suggest multiple windows of functionality for SV2A that need to be dissected further for mechanism and for direct versus indirect effects. Altogether, STAMP provides us with the optimal system needed to explore and discover both known and transient key temporal-spatial mechanisms, setting the stage for more detailed network mapping of the G_0_ program and other biological questions.

## DISCUSSION

This study introduces STAMP, a systematic methodology that employs the power of imaging and phospho-proteomics in a synchronized cell population to identify and dissect detailed changes in cellular PTMs, localization, and function in time and space. We describe and give examples of just some of the many ways that STAMP can be used to determine the sequence of events in multi-step processes (such as ciliogenesis or G_0_ transition). By analyzing cells at different time points after release from mitosis, strong, testable hypotheses emerge. Our stringent synchronization protocol for tracking G_0_ progression enables the enrichment and detection (in many cases for the first time) of rare, transient cellular events and PTMs of low abundant proteins. Note that while we focused herein on phosphorylations, other PTMs may be studied using the same protocols. We describe several applications that have already yielded innovative, exciting directions, though there are clearly many more potential uses for STAMP. Understanding the detailed, transient signaling mechanisms that drive the G_0_ program and ciliary function is pivotal for human health, as disruptions in kinase-substrate binaries have been linked to numerous pathologies. STAMP will allow researchers to determine mechanistically how G_0_ is disrupted in severe pathologies, including cancer, metabolic disease, neurodegeneration, and developmental disorders. We can use this approach to track how specific genetic mutations contribute to defects in ciliary assembly and function in ciliopathies. By elucidating tightly coordinated cellular events, this approach provides a deeper understanding of disease mechanisms and presents a more refined perspective on potential therapeutic targets. This is essential because kinases are involved in numerous signaling pathways, and dissecting their specific roles in distinct events requires precise temporal and spatial resolution. This is particularly crucial for studying G_0_ and distinguishing it from other stages of the cell cycle.

In establishing the timeline using live and fixed cell imaging (Fig.2B), we identified central features of G_0_ and ciliogenesis, including but not limited to a “tubulin barrel” intermediate step lasting from ∼2-4.5hr (which, to our knowledge, has not been previously reported), as well as a hierarchy of recruitment of proteins to specific sites; *e.g*., IFT-B precedes IFT-A to the basal body, and ARL13B is recruited to basal bodies before TTBK2. Each of these observations opens up opportunities for discovering exciting molecular mechanisms. We also demonstrate the use of STAMP to analyze cells deleted for specific components of known pathways (ARL13B, IFT52, and TTBK2) to address and further uncover ordering of processes and their mechanisms. We give examples in which clustering of phospho-signatures, live cell imaging, and chemical inhibitors (PIT-STAMP) can be implemented to generate and begin to test mechanistic hypotheses. For example, we identified key ciliogenesis regulators (featured here, PRKD1 and SV2A (Fig.9-10)), and a novel role for translational control in establishing quiescence. Furthermore, we temporally mapped known and novel phosphorylations to key regulators of NPC assembly (Fig.5) and to autophagy (Fig.6-7) during G_0_ entry. We took this work a step further and tested the hypothesis that we can map specific phosphosignatures of known ciliary proteins to identify key phosphoregulatory steps in essential biological processes. Our discovery that the late-emerging phosphorylation of TTBK2 at residue [S786] fails to rescue ciliogenesis and GPCR recruitment, highlighting new functions for TTBK2 later in the ciliogenesis program (Fig.4). Altogether, these findings solidify STAMP <u>as a means of generating and testing hypotheses concerning the mechanisms driving essential life processes.</u>

Further validation of individual findings is pivotal before drawing hard conclusions. We also acknowledge that the use of chemical inhibitors such as palbociclib and nocodazole for cell synchronization introduces potential artifacts, though we do note here that we have taken numerous steps to control for and mitigate these effects, as outlined in our protocol optimizations and validation experiments. These studies are also limited by temporal resolution, markers used, and processes studied, as our work here is certainly not exhaustive. Here, we focused our studies of the G_0_ program and ciliogenesis in the context of RPEs and MEFs. Based on our early studies, we note differences in the kinetics between RPE and MEF ciliogenesis programs, though the fundamental ordering of the stages of recruitment remains the same. Further work is required to gauge the degree to which these and other cell types vary in their G_0_ programs both on the level of cell function and mechanisms. This work can be extended to other cell lines as well as to more in-depth analyses depending on context-specific questions. Especially interesting would be to extend this work further in the context of already quiescent cells and to perform STAMP in different signaling contexts (*e.g*, SHH pathway, metabolism, neurotransmission, and more).

Overall, we believe that the STAMP method has demonstrated the value of multiplexed studies to provide a more powerful approach to address cell regulation mechanisms. We pioneered an approach that resolves temporally coordinated, transient phosphorylations and cellular events via phospho-proteomics and microscopy, overcoming the typical background signal from differential cell cycle timing that has previously hindered such studies. This is important for dissecting higher-order signaling, or how different processes may be connected in time, space, and mechanism. Other techniques/tools can be multiplexed with STAMP, *e.g.,* phosphosite-specific mutagenesis, RNA-seq, electron and super-resolution microscopy, other PTMs, and other drug screens to name a few. Furthermore, our use of STAMP in both WT and genetically modified RPEs and MEFs supports our conclusion that this approach can be modified for future use in a wide range of cell types, including perhaps cancer cell lines. Indeed, comparisons between cell lines is yet one more way mechanisms and context-specific changes in them are likely to be revealed via STAMP. With STAMP, the ability to map signaling networks that drive biology and to ascertain how lesions in specific mechanisms propagate systemic cellular defects and disease will open doors not only for biology, but also for targeted therapeutic development.

## MATERIALS AND METHODS

### Key notes about this protocol

The primary objective for this protocol was to obtain a pure cell population of cells synchronized at metaphase, not necessarily to achieve high efficiency. We wanted to ensure that the conclusions that we drew from these mass spectrometry and imaging approaches were accurate and to remove the obscurity that comes with having cells in mixed stages of cell cycle. Overall, we have achieved 30% efficiency (*i.e*, if we plated 100 cells, only 30 of them would be synchronized in mitosis by the end of the time course and usable for future assays). Therefore, when planning your experiment, make sure to calculate how many cells you will need at the end and seed at least 5x more cells than you would require. The efficiency issue likely can be resolved with future optimization (i.e. adjusting the timing of Palbociclib treatment and release from G_1_, optimizing drug treatment concentrations). For our purposes, though, we prioritized purity over efficiency to maximize clean, interpretable signals for mass spectrometry. Use of different cell lines will require re-optimization of key steps including incubation times in nocodazole and palbociclib, as well as recovery from drug treatment. This cell-to-cell variability can be attributed to numerous factors, including differential doubling times and rates of cell settling and formation of focal adhesions.

### Materials used

- Cell lines used: WT hTERT RPE-1 originally was obtained from ATCC (CAT) and WT Mouse Embryonic Fibroblast were originally obtained from ATCC (cat #: CRL-2991). All stable cell lines (*i.e*, knockouts, ones expressing fluorescent labels) used in this experiment were generated from the original WT RPEs obtained from ATCC.
- Maintenance medium: DMEM/F12 (Thermofisher; cat #: 12400024) buffered with Sodium Bicarbonate (Gibco; 25080-094) + 10% FBS (Gemini; cat #: 100-106) + GlutaMax (Life Technologies; cat #: 35050-079) + Pen-Strep (Pen-Strep-Glutamine 100x, Gibco; cat #: 10378016)
- Serum starvation medium: DMEM/F12 + GlutaMax + Pen-Strep
- Puromycin: (SIGMA-Aldrich; P9620)
- Blasticidin: (Corning; 30-100-RB)
- CRT 0066101 (dissolved in DMSO): (MedChem Express; HY-15698A)
- Palbociclib (dissolved in DMSO)-2.5mg/mL stocks (Selleck-Chem; cat #: S4482)
- Nocodazole (dissolved in DMSO)-5mg/mL stocks (Cayman; cat #: 13857)
- Phosphate Buffered Saline (made in house using standard lab protocols)
- Appropriate plateware for specific assays:

◦ BioLite 6-well Multidish Cell Culture-treated surface: (Thermo Scientific; cat #: 120184)
◦ BioLite 100mm Tissue Culture Dish: (Thermo Scientific; cat #130182)
◦ Roller bottles InVitro PETG Roller Bottles - 4200cm^2^: (Thermofisher; 12-565-532)
◦ Glass-bottom 96-well plates, black plate with lid (0.17+/− 0.005mm): (Cellvis; P96-1.5H-N)
◦ BioLite 60 mm dish Cell Culture-Treated Surface
- Immunofluorescence reagents:

▪ Microscope coverglass, 12CIR-1.5 (Fisherbrand, 12-545-81)
▪ VistaVision Microscope Slides, Histobond (75×25×1mm) (VWR; cat #: 16005-110)
▪ SlowFade Gold antifade reagent (Invitrogen; cat #: S36936)
▪ DAPI, 10mg/mL (Biotium; cat #: 40043)
▪ Lipofectamine 3000 transfection reagent: (Thermofisher; cat #: L3000015)
▪ OptiMEM: (Thermofisher; cat #: 51985091)
▪ Primary antibodies:

• Acetylated tubulin: (mouse IgG2B; Santa Cruz; Cat #: SC-23950; 1:5000 of 200μg/mL stock)
• Cep170: (mouse IgG1; Thermo Scientific; Cat #: 41-3200; 1:1000 of 0.5mg/mL stock)
• TTBK2: (rabbit; Sigma; Cat #: HPA018113; 1:250 of 0.4mg/mL stock)
• GPR161: (rabbit; Proteintech; Cat #: 13398-1-AP; 1:500 of 0.7mg/mL stock)
• IFT140: (rabbit; Proteintech; Cat #: 174(*18*)60-1-AP; 1:1000 of 0.9mg/mL stock)
• IFT188: (mouse IgG1; Proteintech; Cat #: 60227-1-Ig; 1:1000 of 1.4mg/mL stock)
• Cep290: (rabbit; Abcam; Cat #: ab85728; 1:500 of 1mg/mL stock)
• PDGFRα: (rabbit; Cell Signaling; Cat #: 3174; 1:500 of stock (unknown concentration))
• TULP3: (1:250 of Rabbit polyclonal, made in house)(*21*)
• SV2A: (rabbit; Thermofisher; Cat #: PA5-52476; 1:250 of 0.08 mg/mL)
• PRKD1: (rabbit; Sigma-Aldrich; Cat #: HPA029834; 1:250 of stock (unknown concentration))
• Rab34: (mouse IgG2a; Santa Cruz; Cat #: sc-376710; 1:250 of 0.2mg/mL)
• PCM-1: (rabbit; gift from Dr. Andreas Merdes;(*66*) 1:1,000,000 of stock)
• ARL13B: (mouse IgG1; Santa Cruz; Cat #: sc-515784; 1:1000 of 0.2mg/mL stock)
• ARL13B: (rabbit; Proteintech; Cat #: 17711-1-AP; 1:1000 of 0.9mg/mL)
• FOP: (mouse IgG2B; Abnova; Cat #: H00011116-M01; 1:1000 of 0.1mg/mL)
• GM130: (rabbit; Cell Signaling; Cat #: 12480T; 1:000 of stock (unknown concentration))
• TGN38: (mouse IgG1; BD Biosciences; Cat #: 610898; 1:1000 of 0.25mg/mL)
• LAMP1: (rabbit; Abcam; Cat #: ab24170; 1:1000 of 1mg/mL)
• HA.11 epitope: (mouse IgG1; BD Biosciences; Cat #: 90150; 1:1000 of 1mg/mL)
• IFT172: (mouse IgG1; Santa Cruz; Cat #: sc-398393; 1:500 of 0.2mg/mL)
• IFT20:(rabbit; Proteintech; Cat #: 13615-1-AP; 1:500 of 0.4mg/mL)
• IFT43: (rabbit; Sigma; Cat #: HPA003438; 1:500 of 1mg/mL)
▪ Secondary antibodies: Alexafluors from Invitrogen

• Donkey-anti-Rabbit 488 (cat #: A-32790)
• Donkey-anti-Rabbit 568 (cat #: A-10042)
• Donkey-anti-Rabbit 647 (cat #: A-21447)
• Goat-anti-mouse IgG1 488 (cat #: A-21121)
• Goat-anti-mouse IgG1 568 (cat #: A-21124)
• Goat-anti-mouse IgG1 647 (cat #: A-21240)
• Goat-anti-mouse IgG2a 488 (cat #: A-21131)
• Goat-anti-mouse IgG2a 568 (cat #: A-21134)
• Goat-anti-mouse IgG2a 647 (cat #: A-21241)
• Goat-anti-mouse IgG2b 488 (cat #: A-21141)
• Goat-anti-mouse IgG2b 568 (cat #: A-21144)
• Goat-anti-mouse IgG2b 647 (cat #: A-21242)
▪ Paraformaldehyde 16% Solution EM grade 10X10 ML (Electron Microscopy Sciences; cat #: 100503-917)
▪ Methanol (Fisher Scientific; cat #: A412-4)
▪ Triton X-100 (Fisher Scientific; cat #: AC21568-2500)
▪ NP-40 substitute (Roche Applied Sciences; cat #: 11754599001)
- Phospho-proteomics reagents

◦ Sodium deoxycholate (Sigma; cat #: D6750)
◦ 2-Chloroacetamide (Sigma; cat #: C0267)
◦ Potassium hydroxide (Sigma; cat #: P5958)
◦ Calcium chloride (Sigma; cat #: 793639)
◦ Trifluoroacetic acid (Fisher; cat. #: AAL06374AC)
◦ Isopropanol (Fisher; cat. #: A461-1)
◦ Acetic acid (Fisher; cat. no. A11350)
◦ Tris(hydroxymethyl)aminomethane hydrochloride (Tris-HCl; cat. No. AC228031000)
◦ LC/MS Grade (Optima TM; cat. #: W6-4)
◦ Formic acid, 99.5%, LC/MS Grade (Optima TM; cat. #: A117-50)
◦ Tris(2-carboxyethyl)phosphine hydrochloride (Thermo Fisher; cat. #: PG82080)
◦ PierceTM BCA Protein Assay Kit (cat. #: 23225)
◦ PierceTM Bovine Serum Albumin (BCA) Standard Pre-Diluted Set (cat #: 23208)
◦ PierceTM Quantitative Colorimetric Peptide Assay (cat # 23275)
◦ Ammonia solution, 25% wt/vol (Honeywell; cat. #: 44273)
◦ Acetonitrile LC/MS grade (Honeywell; cat. #: 14261-1L)
◦ Potassium dihydrogen phosphate (Merck; cat. #: 5438410100)
◦ Methanol LC/MS grade (Millipore Sigma; cat. #: 900688-1L)
◦ Trypsin/Lys-C Mix, Mass Spec Grade (Promega, cat. #: V5073)
◦ Empore C18 47 mm Extraction Disk (Empore, cat. #: 320907D)
◦ Halt™ Protease Inhibitor Cocktail, EDTA-Free (Thermo Scientific, cat. #: 78425)
◦ Halt™ Phosphatase Inhibitor Cocktail, EDTA-Free (Thermo Scientific, cat. #: 78428)
◦ Bruker PepSep C18 10cm packed 1.5 μm beads in 150 μm I.D column (Bruker; cat. #: 1893483) attached to the ZDV Sprayer (I.D. 20 μm, cat. #: 1865710)
- Live cell imaging reagents

◦ SiR Tubulin live cell imaging dye (Cytoskeleton, Inc.; cat #: CY-SC002)
◦ Imaging medium (phenol-red free, supplemented with HEPES, serum free) (Thermofisher; cat #: 11039021)
◦ HEPES, 1M (Gibco; cat #: 15630080)
◦ MatTEK glass bottom plates (MatTEK; cat #: P35G-1.5-14-C)
◦ IBIDI 8-well glass bottom chamber slides (IBIDI, Cat #: 80827)
- Drug inhibitor treatments

◦ SV2A inhibitor (levetiracetam) (Millipore-Sigma; cat #: L8668-50MG)
◦ Cycloheximide (Sigma-Aldrich; cat #: C7698-1G)
◦ Dynarrestin (Sigma-Aldrich; cat #: SML2332-5MG)
◦ Dimethyl sulfide (DMSO) anhydrous, ≥99.9% (Sigma-Aldrich; cat #:276855)

## GENERATION AND MAINTENANCE OF STABLE CELL LINES

hTERT RPE-1 cells (ATCC; Cat #: CRL4000), HEK-293 (ATCC; Cat #: CRL-1573), as well as all stable cell lines generated from this mother population, were grown in DMEM/F-12 (12400024, Thermo Fisher Scientific) supplemented with 10% FBS (100–106, Gemini), 1× GlutaMax (35050-079, Thermo Fisher Scientific) at 37°C in 5% CO2. To induce cilium formation, RPE cells were incubated in DMEM/F12 supplemented with 1×GlutaMax (serum-free media). Cell lines were authenticated via a short-tandem-repeat based test. Mycoplasma negativity was confirmed via DAPI staining for all immunofluorescence experiments, screening for mycoplasma of all cell lines at least every month. All cells were maintained below passage 15 to prevent phenotypic/genetic drift and were raised without antibiotics except for endpoint assays to prevent the propagation of contaminated stocks.

CRISPR KO cell lines were generated as previously described.(*20*)

Stable cell lines expressing fluorescently tagged constructs were generated using lentivirus, as previously described(*20*). Lentivirus carrying either gene of interest was produced by co-transfecting HEK293 cells with 150 ng of pCMV-VSV-G, 350ng of pCMV-dR8.2 dvpr, and 500 ng of lenti-viral transfer plasmids (previously described) along with 3μL of Fugene 6 (E2692, Promega) transfection reagent. Medium was replaced 24hr after transfection to prevent toxicity from overexposure to transfection reagent, and virus was harvested at 48 hr post-transfection. Virus was then filtered with a 0.45 μm PVDF filter (SLHV013SL, Millipore) and mixed with 4-fold volume of fresh media containing 12.5μg/mL polybrene (TR-1003-G, Millipore). Following infection for 66hr, cells were selected with 10μg/mL blasticidin (30-100-RB, Corning) for at least 10 days before subsequent analysis.

## PLASMIDS/CLONING

TTBK2 entry gene was purchased from DNASU. The [S786A] mutant was generated by PCR using Site-Directed Mutagenesis (Thermo Fischer Scientific Catalog # F541) and appropriate mutagenesis primers. Both the WT-TTBK2 and S786A-TTBK2 were introduced into destination pCS2+HA-tagged expression vectors by Gateway cloning (Thermo Fischer Scientific11791020). Plasmids were sequenced to verify the integrity of recombination and mutation status by whole plasmid sequencing (Elim Bio).

### Primers used

TTBK2_S786A_Reverse: TAGAGGCGGATAATGAAGATGAGAAGTTAAGT

TTBK2_S786A_Forward: CTTCATTATCCGCCTCTAAAAGGATGCTTTTCTCTTC

## TRANSIENT TRANSFECTION

TTBK2 KO RPE cells were transiently transfected for ciliogenesis rescue experiments using Lipofectamine 3000. In brief, cells were plated onto glass coverslips in the bottom of 6-well plates at 80% confluence, 24hr before transfection. The day of the experiment, we swapped the medium for each well with 2mL of Opti-MEM transfection medium. Plasmids were transfected into cells for 4 hours, using 2.5μg DNA + 5uL P3000 reagent + 10uL lipofectamine per well (with the complexed plasmid being diluted in 250μL OptiMEM before being pipetted dropwise onto the wells). After the transfection, the wells were washed 2x with PBS before medium was swapped to serum-free DMEM/F12 + GlutaMax. Cells were serum starved for 24hr before being fixed and stained for IF.

## CELL SYNCHRONIZATION PROCEDURE

1. RPE cells were seeded at 10% confluence to grow up for 24 hours (allowing them to double) or seeded at 20% confluence. Ensure that the cells are evenly dispersed over the plate and avoid having any clumps. Cells that clump together tend to be less responsive to cell synchrony treatment, leading to decreased yield and/or purity.

2. 24hrs later, spike in Palbociclib at a final concentration of 1μg/mL. Incubate for 18hr at 37°C. This will arrest the cells in G_1_.

***NOTE:*** *Since the start of this project, we have observed that <u>reducing the incubation time from 18hr to 15hr greatly increases the yield of synchronized cells</u> (reaching ∼40-50% yield). We would highly recommend others to use 15hr palbociclib treatment as their starting point and to continue to optimize from there if seeking to improve yield*.

3. Aspirate medium and wash at least 1x with PBS. Add fresh maintenance medium and let cells release from G_1_ for 8hr.

4. Aspirate medium and replace with maintenance medium supplemented with nocodazole at a final concentration of 50ng/mL. Incubate at 37°C for 12 hr.

5. Check the efficiency of synchronization via brightfield microscopy before proceeding. If successful, you should see several rounded-up cells floating in the medium or still attached to the plate. These rounded cells are blocked in mitosis at approximately metaphase. At this time there will also be some cells that are still flat and attached to the plate. It is pivotal to only harvest the cells in mitosis and to avoid collecting the adherent cells.

6. Perform mitotic shakeoff-firmly tap on the side of the plates using the palm of your hand to try to increase detachment of rounded, mitotic cells from the plate. Use a pipettor to aspirate the supernatant and use it to ***<u>gently</u>*** wash off the mitotic cells. DO NOT be too rigorous, and DO NOT scrape the bottom of the plate or vortex the cells. This step is pivotal for ensuring the purity of your sample and to avoid contamination with cells from other cell cycle stages.

7. Spin down cells at 500rpm for 5 min and aspirate supernatant. Resuspend in 10mL PBS to wash away remaining nocodazole medium.

8. Spin down cells at 500rpm for 5 min and aspirate supernatant. Resuspend in 10mL maintenance medium (with 10% FBS) and incubate the cells for 20 minutes at 37°C, KEEPING THE CELLS IN THE CONICAL TUBE. This step is pivotal-incubating the cells for a short period in maintenance medium before thrusting them into serum starvation and before plating the cells allows the cells to recover faster and ciliate faster. Plating cells directly into serum starvation medium without including a recovery step increased the time for axonemogenesis to occur by 6-8hr. We suspect it has something to do with helping to replenish the soluble pool of tubulin, though this would require further testing.

9. Spin down the cells at 500rpm for 5 min, aspirate supernatant and resuspend in serum starvation medium. Make sure that the cells are not clumpy and have no signs of contamination with adherent cells (adherent cells that accidentally got scraped off tend to look less round and symmetric compared to mitotic cells) using visual inspection with brightfield microscopy. We also do the added precautionary step of passing the cells through a single-cell filter to avoid clumps that could contaminate the sample. Count the cells, and aliquot into appropriate vessels depending upon the purpose of your experiment.

10. When synchronizing cells, we also take into account other variables that may impinge on the reproducibility of time windows. Make sure to perform all seeding quickly and efficiently from start to finish to avoid the added variable of time for adherence. Place each individual plate on the shelf of the incubator rather than stacking plates to maintain as constant a temperature as possible between samples. Use fibronectin-coated (10µg/mL) vessels for all experiment described below, to ensure that the cells still adhere even in the absence of serum. Avoid taking cells in and out of incubator except during harvesting to limit potential effects of temperature on G_0_ program entry.

## IMMUNOFLUORESCENCE PROCEDURE

1. For immunofluorescence, cells were plated onto acid-washed, fibronectin-coated coverslips inside 35mm individual dishes. This allows for ready harvest of multiple individual time points. Metaphase-synchronized cells were seeded at 50-60% confluence to ensure that there are plenty of cells to make that first doubling (approximately 5-6×10^5 cells per 35mm plate, per time point). Make sure to add the cells dropwise to fibronectin-coated coverslips and plates filled with serum free medium rocking back and forth to ensure that the cells are evenly distributed.

2. Place plates in the incubator at 37°C, making sure not to stack plates but to put each separately on the rack. This is to ensure that they all reach the same temperature at the same time, which is critical for microtubule dynamics and thus synchronicity of ciliogenesis/ G_0_ progression.

3. Collect time points, fixing and staining cells according to standard in-house protocols. Some of the following preliminary markers to ensure the success of the synchronization protocol:

a. Acetylated tubulin- universal marker of ciliary axoneme (some markers like ARL13B do not show up in every cilium depending on the genotype). I would always recommend having an acetylated tubulin channel on your coverslip for all comparisons, if possible.

i. Another bonus is that acetylated tubulin also serves as a marker for the midbody and other key features in mitosis, so this coupled with a nice DAPI/nuclear stain can help act as a quality control check to make sure you really synchronized your cells.
b. Cep170, centrin, or gamma tubulin to mark the basal body-this will help give you some sort of framework to tell if certain ciliogenesis markers have recruited/disappeared.
c. Cep164 and TTBK2 as early markers of ciliogenesis
d. Cep290 and NPHP4 for transition zone
e. RAB34 for vesicle recruitment
f. GPR161 and PDGFR-alpha to mark ciliary receptors.
g. IFT88, IFT20, IFT172 (IFT-B), IFT140, IFT43 (IFT-A), and TULP3 to mark ciliary traffic machinery.

4. When performing mass spectrometry or any other experiment, collect time points for IF simultaneously. If using 10cm plates (e.g, for mass spectrometry), place a coverslip in the 10cm plate to directly correlate cellular phenotypes with molecular readout. Be aware that while some fixed coverslip samples can be stored long term at 4°C like in the case of PFA fixation, methanol fixed samples do deteriorate 24hrs after fixation. In this case, it is advised to perform IF the same day as sample collection.

## PHOSPHO-PROTEOMICS PROCEDURE

Stock solutions were prepared for the experiments, including 1 M Tris-HCl (pH 8.5), 5M potassium hydroxide (KOH), 100 mM KH2PO4, and 2 M calcium chloride (CaCl2). All stock solutions were stored at room temperature (RT; 20–25°C).

10cm plates of cells (4×10^6 metaphase synchronized cells, accounting for the cells doubling to 8×10^6 cells) were collected for each replicate for each time point and processed for phospho-proteomics.

SDC lysis buffer was formulated with 4% (wt/vol) SDC, 100 mM Tris-HCl (pH 8.5), 1x Halt™ protease, and phosphatase Inhibitor cocktail. It is crucial to prepare this buffer freshly as SDC crystallizes upon storage in solution.

Reduction/alkylation buffer was composed of 100 mM TCEP and 400 mM 2-chloroacetamide (CAM), with pH adjusted to 7–8 using KOH. The pH of the buffer was verified using a pH indicator strip. This buffer had to be prepared immediately before use to ensure the full activity of CAM, and any excess had to be safely discarded after use.

Trypsin buffer was prepared by combining 0.05% (vol/vol) AcOH and 2 mM CaCl2. This buffer was stored at −20°C, where it can remain stable for more than one year. For reconstitution, 1 mg of lyophilized trypsin/Lys C was resuspended in 1mL of trypsin buffer (1 mg/mL). The enzymes were resuspended by vortexing and then were then centrifuged (1,000g for 1 min at RT). Phosphoproteomic loading buffer was formulated with 6% (vol/vol) TFA/80% (vol/vol) ACN. Phosphoproteomic enrichment buffer consisted of 48% (vol/vol) TFA and 8mM KH_2_PO_4_.

Wash buffer was prepared using 5% (vol/vol) TFA/60% (vol/vol) ISO. Elution buffer was prepared by adding 200 µL of ammonia solution (NH4OH) to 800µL of 40% (vol/vol) ACN.

The SDC lysis buffer was chilled to 4°C. A 300 µL volume of lysis buffer was added to 1×10^6 cells to achieve a protein concentration of approximately 3 mg/mL. Lysates were immediately heat-treated for 5 min at 95°C to facilitate lysis and inactivate endogenous proteases and phosphatases. Then, lysates were homogenized by sonication at 4°C. Disulfide bonds and carbamidomethylated cysteine residues were reduced by adding a 1:10 volume (30 µL) of reduction/alkylation buffer to the samples. Samples were incubated for 5 minutes at 45°C with shaking at 1,500rpm. After removing the samples from heat and allowing them to cool to room temperature (RT), Lys-C and trypsin enzymes were added at an enzyme-to-substrate ratio of 1:100 (wt/wt), and the samples were digested overnight at 37°C with shaking at 1,500 rpm. To each sample, 400µL of ISO was added and thoroughly mixed at 1,500 rpm for 30s. Proper mixing with ISO was ensured before proceeding to the addition of EP enrichment buffer to prevent precipitate formation. A 100µL volume of EP enrichment buffer was added to the samples, and the samples were mixed thoroughly at 1,500 rpm for 30 seconds. The samples were carefully inspected for the presence of precipitate or cloudiness. If any precipitate was observed, the samples were cleared by centrifugation (2,000g for 15 min at RT). The supernatants were then carefully transferred to clean tubes before TiO2 beads were added. The TiO2 beads were resuspended in EP loading buffer at a concentration of 1mg/µL. An aliquot of 20mg suspended beads was pipetted into each sample, then incubated at 40°C with shaking at 2,000rpm for 5 min. The beads were pelleted by centrifugation (2,000g for 1 min at RT), and the non-phosphopeptide supernatant was discarded using a glass aspirator tip attached to a vacuum hose. 1mL of EP wash buffer was added to the samples. Samples were then incubated at RT with shaking (2,000rpm) for 30s. The beads were pelleted by centrifugation (2,000g for 1 min at RT). The supernatant was then discarded. Washing was repeated four more times, and the same wash procedure was performed. After the final wash, the beads were resuspended in 75µL of EP.

Homemade Stage-Tips were constructed using two C18 Empore disks, following established procedures. The fabricated Stage-Tips were washed two times with 100μL of methanol, one time with 100μL of 80% acetonitrile/0.1% acetic acid, and two times with 100μL of 1% acetic acid. Enriched phosphopeptides were loaded onto the Stage-Tips in 100μL of 1% acetic acid. Subsequently, the Stage-Tips were washed three times with 100μL of 1% acetic acid to remove salts. Finally, the phosphopeptides were eluted from the Stage-Tips using two elution steps of 30μL each, with 80% acetonitrile/0.1% acetic acid as the elution buffer.

### Liquid Chromatography Setup

A nanoELute ultra-high-pressure nano-flow chromatography system was used and directly coupled online with a hybrid trapped ion mobility spectrometry—quadrupole time-of-flight mass spectrometer (timsTOF Pro, Bruker) employing a nano-electrospray ion source (CaptiveSpray, Bruker Daltonics).

### Chromatographic Conditions

The liquid chromatography was conducted at a constant temperature of 50°C, employing a reversed-phase column (PepSep column, 10 cm × 150 μm i.d., packed with 1.5 μm C18-coated porous silica beads, Bruker) connected to the 10 μm emitter (Bruker). The mobile phase consisted of two components: Mobile Phase A, comprising water with 0.1/2% formic acid/ACN (v/v), and Mobile Phase B, comprising ACN with 0.1% formic acid (v/v).

### Gradient Elution

Peptide separation was achieved using a linear gradient from 2-33% Mobile Phase B within 60 min. This was followed by a washing step with 95% Mobile Phase B and subsequent re-equilibration. The chromatographic process maintained the flow rate at 400nL/min.

### MS Acquisition

Samples were analyzed using the timsTOF HT Mass Spectrometer in DDA-PASEF mode. The TIMS elution voltage was calibrated linearly to obtain reduced ion mobility coefficients (1/K0) by using three selected ions from the Agilent ESI-L Tuning Mix (m/z 622, 922, 1222). The mass and ion mobility ranges were set from 100 to 1700 m/z and 0.7 to 1.3 1/K0, respectively. Both ramp and acquisition times were set at 100 ms. Precursor ions suitable for PASEF-MS/MS were chosen from TIMS-MS survey scans using the PASEF scheduling algorithm. A polygon filter was applied to the m/z and ion mobility plane to prioritize features likely representing peptide precursors over singly-charged background ions. The quadrupole isolation width was set to 2 Th for m/z < 700 and 3 Th for m/z > 700, with collision energy linearly increased from 20 to 60 eV as ion mobility ranged from 0.6 to 1.6 (1/K0).

### Data Analysis

Raw data files were processed using MS Fragger software against the NCBI Homo sapiens RefSeq protein database. Search parameters included CID fragmentation with a precursor error tolerance of 20 ppm and a fragment ion tolerance of 40 ppm. Searches included S/T/Y phosphorylation and up to three modifications per peptide besides standard modifications. Peptides were validated using Percolator and Protein Prophet at 1% FDR. Protein quantification was performed using IonQuant, with normalization across runs and Match Between Runs settings accommodating retention time and ion mobility tolerances of 0.4 minutes and 0.05 (1/K0), respectively. (*67*)

To create the heatmap, we first extracted the proteins and their identified phosphorylation sites and established a series of defined time points that represent the intervals at which measurements were taken. We calculated average intensity values, accounting for any missing data. The intensity values for each phosphorylation site were normalized by dividing each intensity value at any given time point by the average intensity across the time for the same phosphorylation event. The putative kinases were assigned based on the Kinase prediction tool available on Phosphosite. Once normalized, hierarchical clustering was then performed, where correlations between observations were computed to determine dissimilarities between the phosphorylation sites based on their intensity patterns over time. This clustering allowed us to group similar phosphorylation events and kinases based on their response patterns across the timepoints. The number of clusters was optimized by setting a minimum threshold or cutting the dendrogram to ensure an appropriate number of distinct clusters. Finally, a heatmap was generated to visually represent these clusters, with annotations and a color palette highlighting the relationships between the grouped elements, facilitating the interpretation of the intensity changes across the different conditions. Using the comprehensive kinase substrate specificity profiles established by Cantley’s lab, (*52*) we modified the scoring system to enable more accurate Position-Specific Scoring Matrices (PSSMs). This was achieved by incorporating amino acid frequency distributions from the human proteome. While Positional Scanning Peptide Array (PSPA) data reflects amino acid preferences under controlled conditions in Cantley’s algorithm, incorporating frequencies of amino acids relative to the phosphorylation sites in the human database is essential for better estimation of scores for putative kinases and as a consequence their ranking. For example, if two amino acids exhibit the same favorability in PSPA results but one is less frequent in the specified position relative to the phosphorylation site, it should increase the calculated score compared to the more frequent one. The modified scoring system, which accounts for amino acid frequency, is as follows, where the values of the adjusted corresponding amino acids in specific positions based on their frequencies are multiplied, and then scaled by the probability of a random peptide:

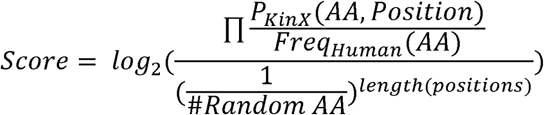

The final kinase rankings were determined by calculating the percentile position of each kinase’s score within the distribution of scores generated from analyzing all available serine/threonine phosphorylation sites from PhosphoSite.(*1*) The phosphorylation signaling maps are only plotted for the highly confident phosphorylations sites, manually inspected by Skyline. MS2 scans are extracted and confirmed for each phosphorylation site demarcated in the figures (Supplementary Data 5).^45^

## LIVE CELL IMAGING PROCEDURE

1. Precoat glass bottom, live cell imaging plates (Ibidi 8-well; Cat #: 80827) for 30 minutes at 37°C with fibronectin (Sigma; FC010-5MG) diluted in PBS (10μg/mL)

2. Plate cells at ∼60% confluence directly into serum free imaging medium in live cell imaging plate (phenol red free DMEM/F12 + Glutmax, supplemented with HEPES), anticipating at least a doubling to get 90-100% confluence to ensure optimal conditions for inducing ciliogenesis. For the purposes of this study, we used cells stably expressing RABL2-GFP and IFT88-mCherry to visualize ciliogenesis. We incubated the cells with SiR-Tubulin live cell imaging dye (far red) at 1:500 dilution to visualize axonemogenesis. Different live cell imaging markers can be used to address different experimental questions using the same protocols.

3. Place cells on stage of live-cell imaging chamber-equipped confocal. We used an Inverted Zeiss LSM 880 Laser Scanning Confocal Microscope. Z-projections (10-12 slices each) were collected either every 8 minutes from 4 different fields over a 24hr time course or every 2 minutes for 1 field. These conditions can be optimized based on the question, including increasing the number of time points, reducing the number of slices, focusing on specific time windows rather than the full ciliogenesis time course, etc.

4. **<u>Tips for reducing phototoxicity:</u>**

a. Increasing the medium in the well helps to reduce the concentration of ROS produced due to light exposure.
b. Maintain a temperature (37°C) and CO_2_ (5%) controlled environment throughout the procedure.
c. Optimize the number of time points and z-stacks collected per well to increase information without risking damaging the cells. Depending on the nature of your question, you may want to prioritize cell health and endurance over detailed temporal/spatial resolution.
d. Minimize laser power and gain. Data processing by adjusting the lookup tables can help with letting you see your data without risking cell damage.
e. Live cell imaging medium supplemented with 10mM HEPES (to modulate pH) and that is phenol-red free (to reduce autofluorescence) helps reduce phototoxicity.

## QUANTIFICATION AND STATISTICAL ANALYSIS

Unless otherwise stated, 100 cells were scored per each replicate, and all experiments were performed in at least biological triplicate and scored in at least biological duplicate. Data were processed using Excel and graphed using GraphPad Prism, and error bars shown indicate the SEM for the data set. Individual data points signify the average of technical replicates for each individual cell line. Statistical significance of the difference between individual test groups was assessed using either One-Way or Two-Way ANOVAs for conditions in which 3 or more samples are being compared, or Student’s t-test for conditions where only 2 samples are being compared: * = p < 0.05; ** = p < 0.01; *** = p < 0.001, **** = p < 0.0001.

## ACKNOWLEDGEMENTS

This work was made possible by the Stanford Cell Sciences Imaging Facility. We would like to extend special thanks to Tomoharu Kanie from UOHSC, Michael East from UNC Chapel Hill, and Richard Kahn from Emory University for their critical feedback and input, as well as to Israel Larios and Roy Ng for providing instrumental technical support throughout the detailed experimental procedures.

## Author Contributions

Conceptualization: MOA, RET, PKJ

Methodology: MOA, RET, AA, LAX

Software: MOA, RET, MM, KYLG

Validation: MOA, RET, AA, KYLG

Formal Analysis: MOA, RET, AA, LAX, KYLG, MM

Investigation: MOA, RET, AA, LEL, LAX, KYLG, PKJ

Resources: PKJ

Data curation: MOA, RET, KYLG

Writing-original draft: MOA, RET, KYLG, PKJ

Writing-reviewing and editing: MOA, RET, KYLG, PKJ

Visualization: MOA, RET, KYLG,

Supervision: MOA, RET, PKJ

Project Administration: PKJ

Funding Acquisition: MOA, RET, AA, PKJ

## Data and materials availability

All data needed to evaluate the conclusions in the paper are present in the paper and/or the Supplementary Materials. Cell lines and plasmids will be made available pending scientific review and a completed material transfer agreement. Requests for cell lines and plasmids should be submitted to.

