## supplemental data for "Synchronized Temporal-spatial Analysis via Microscopy and Phospho-proteomics (STAMP) of Quiescence"

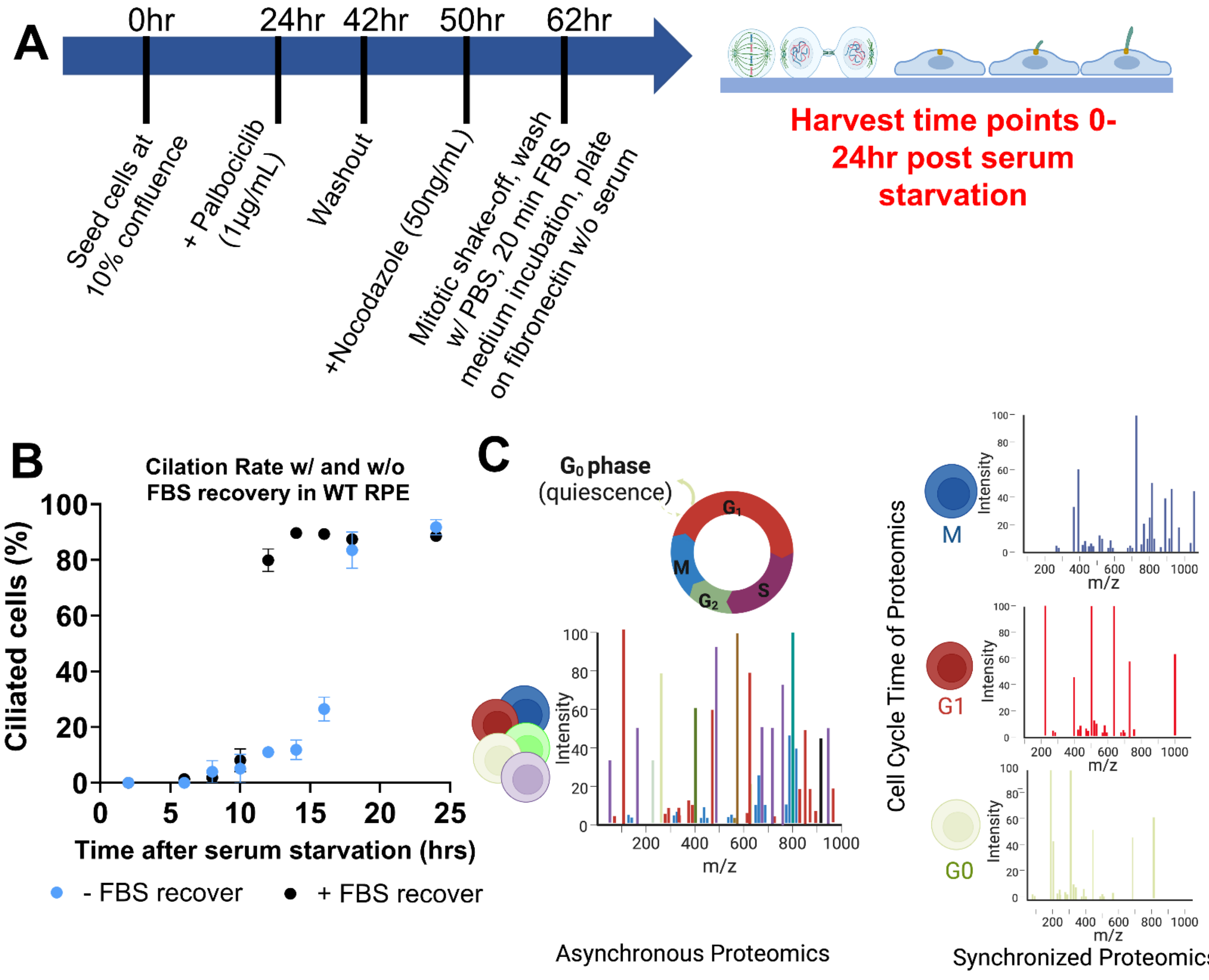


**Fig. S1. Workflow of the STAMP synchronization protocol.** (A) RPE and MEF cells were seeded at low density and were exposed to multiple different rounds of synchronization procedures before being harvested via mitotic shake-off. Schematic of the different stages captured by our synchronization protocol: metaphase onward through ciliogenesis and $G_0$ entry. (B) Quantification of ciliation of FBS-recovered versus non-FBS-recovered STAMP synchronized cells as a control for synchronization protocol. Experiments were performed in replicate, N=100 per replicate. Error bars = SEM. (C) Schematic illustrating the proof of principle for generating synchronized proteomics data, which incorporates the cellular time dimension and enriches for specific proteoforms that are either not detected or quantifiable in asynchronous proteomics.

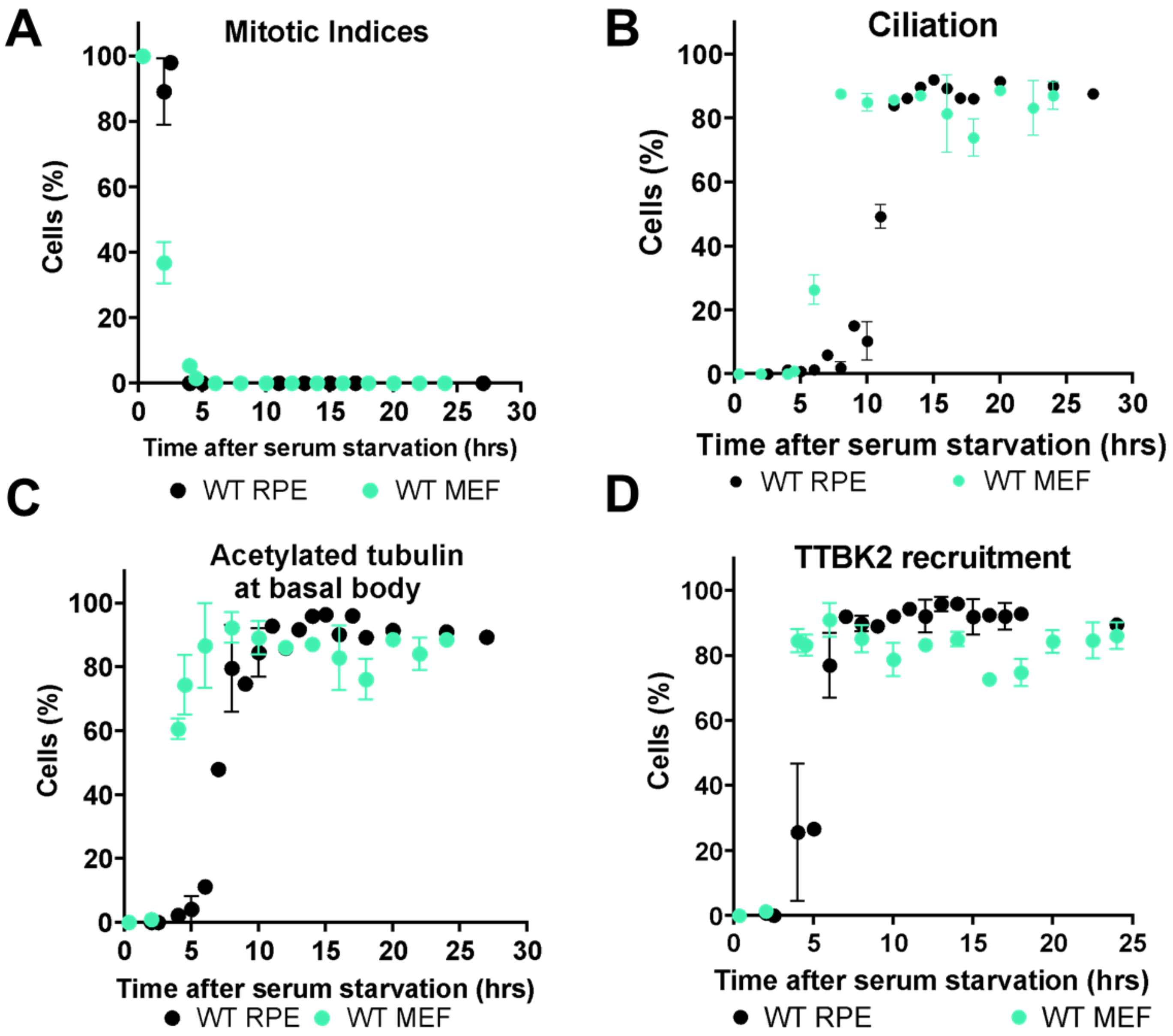


**Fig. S2**. Comparative STAMP of MEFs versus RPEs. Quantification of key markers of ciliogenesis/$G_0$ entry in RPEs versus MEFs via IF, including (A) mitotic indices, (B) ciliation, (C) acetylated tubulin at basal body, and (D) TTBK2 recruitment. Experiments were performed in replicate, N=100 per replicate. Error bars= SEM.

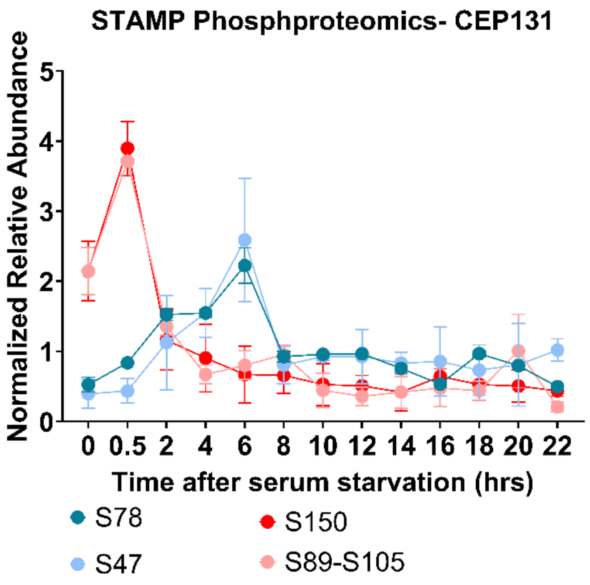


**Fig. S3. STAMP phospho-proteomics identifies transient CEP131 phosphorylation sites.** Plot of the normalized relative abundance of different CEP131 phosphorylations during $G_0$ progression. While proline-directed sites including [S150] and [S89-S105] are much more abundant during cell division, the N-terminal sites ([S47], and [S78]) are dynamically regulated during vesicle recruitment. Error bars = SD.

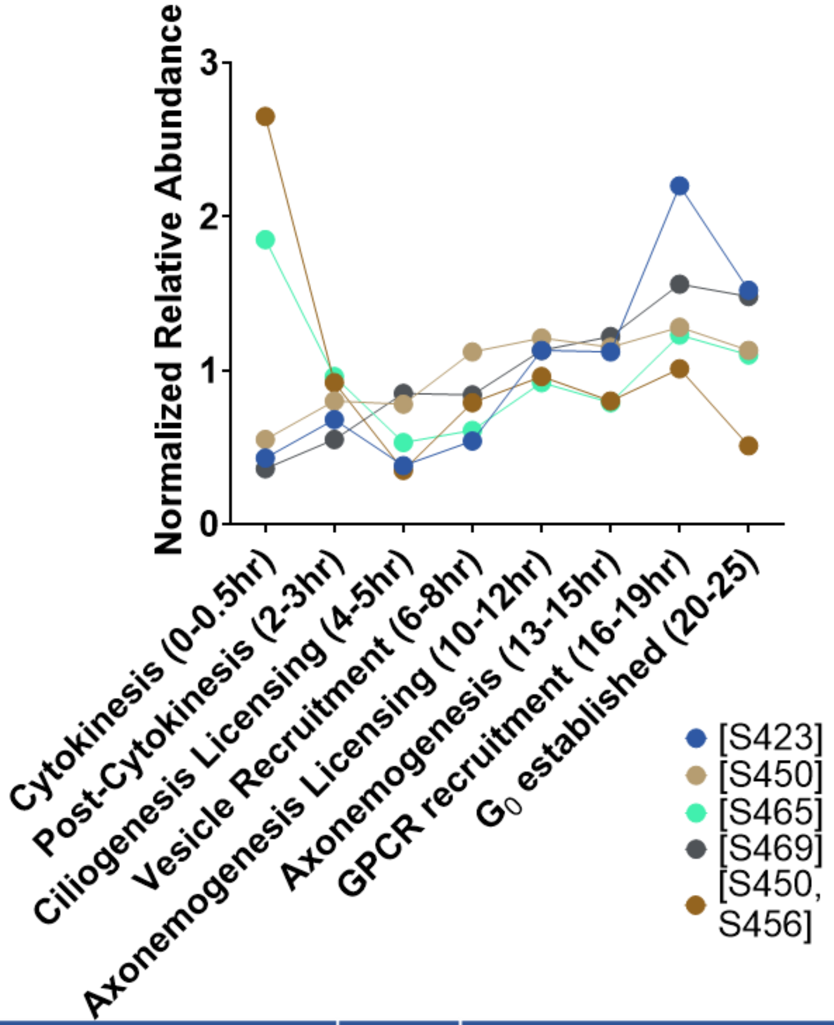


| Site | Motif | Cluster | Predicted window of fxn | Putative Kinase |
|---|---|---|---|---|
| S423 | GRAGPFS**s**SRCGASV | 16 | GPCR recruitment and loading | PKCA |
| S450 | RIERNLQ**s**PTQFQTP | 2 | gradual increase starting at 4,5hr | JNK |
| S450,S456 | RIERNLQ**s**PTQFQ**t**P | 7 | high during cytokinesis | ERK |
| S465 | RSSAIRR**s**GSTSPLG | 7 | high during cytokinesis | PAK |
| S469 | IRRSGST**s**PLGFARA | 2 | gradual increase starting at 4,5hr; Known to induce protein degradation | PBK |

**Fig. S4. Representative example of how to read phosphosignature maps using ULK1.** Above, example plot of the average normalized intensity of 5 different phosphosites of ULK1 over time as generated via STAMP phosphoproteomics. Below, a chart indicating the phosphosite, its corresponding motif, putative kinases that are strong candidates for making that modification (based on the Cantley Kinase Metric), and predicted function.

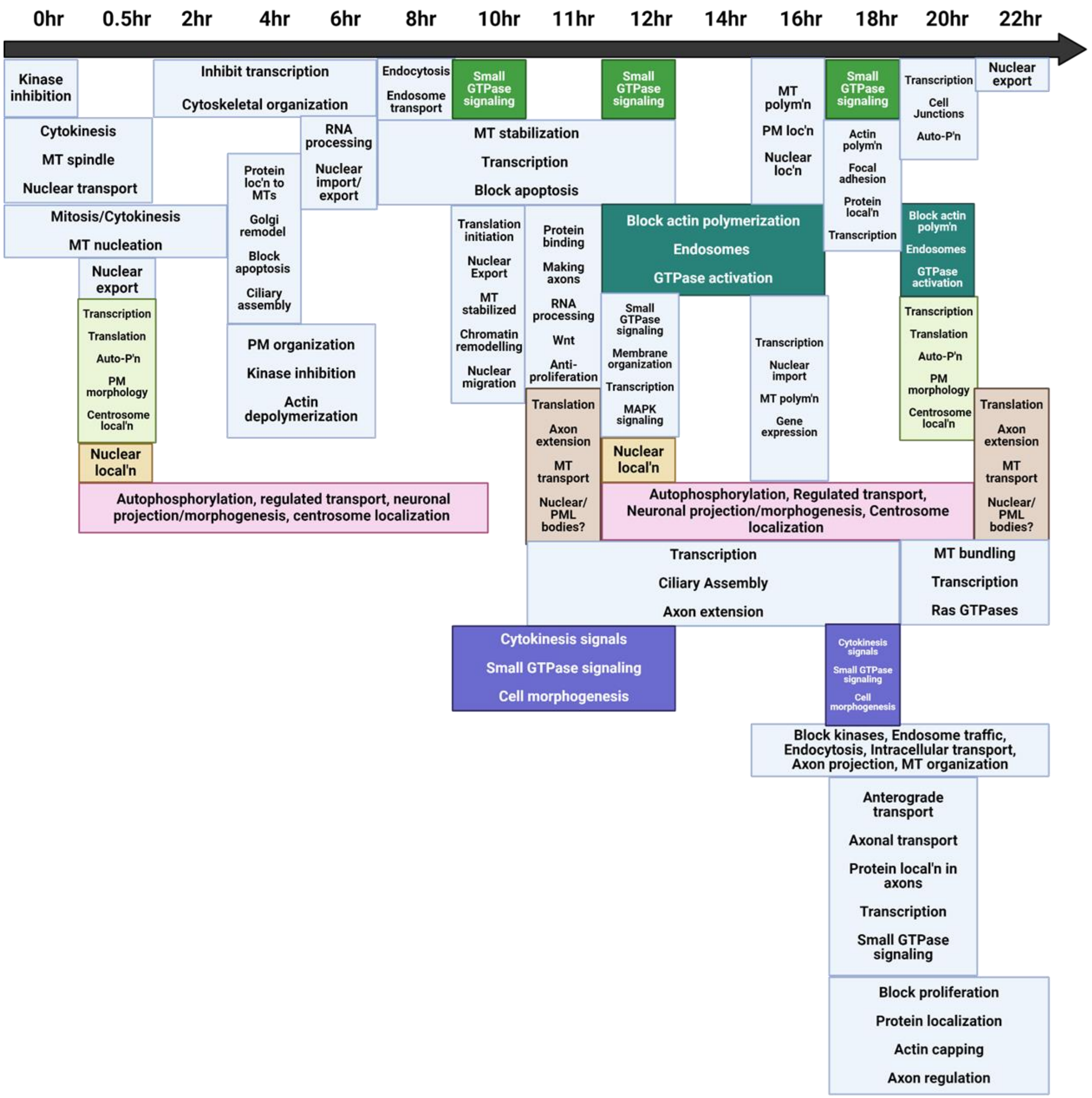


**Fig. S5. Gene Ontology Mapping. Schematic summarizing the Gene Ontology analysis via EnrichR** (https://maayanlab.cloud/Enrichr/). Individual clusters generated from phospho-proteomics (see Fig.5) were run through the software, compared with the GO Biological Process 2023 analysis, and plotted via Bar Graphs and Clustergram. The top 4-5 functions for each cluster were mapped to the timeline as an unbiased approach to observe what functions are active at different time periods during $G_0$ entry.

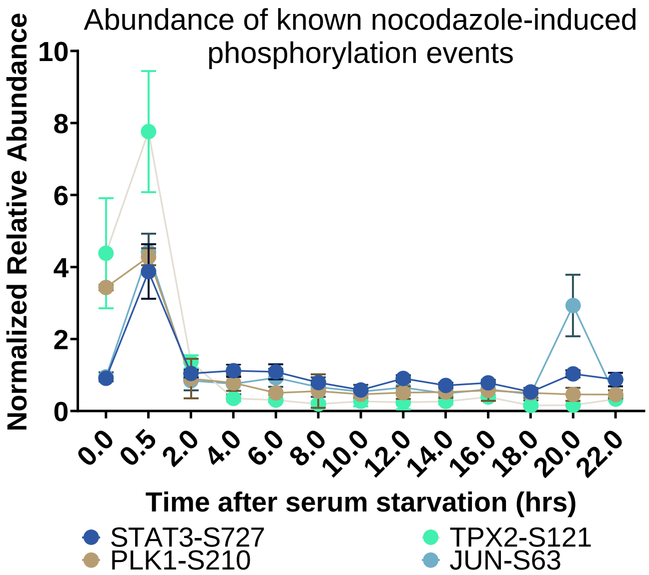


**Fig. S6. Abundance of known nocodazole-induced phosphorylations.** Plot of the intensities of 4 different phosphosites over time that previous groups have shown are signatures of nocodazole treatment, as generated from STAMP phosphoproteomics. Error bars = SD.

### STAMP for “Tubulin Barrel” Disassembly and IFT88 dispersion

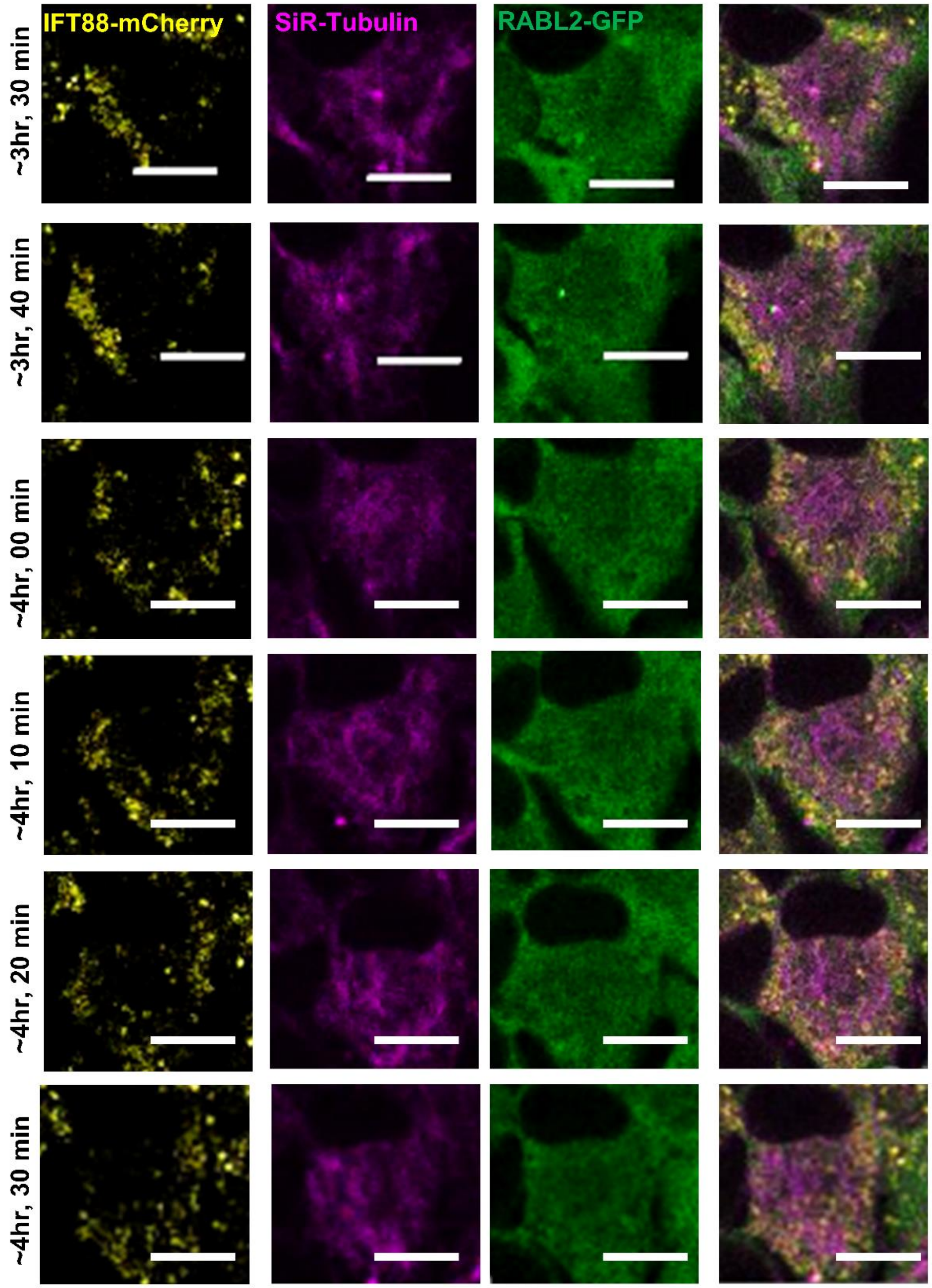


**Fig. S7. STAMP for “Tubulin Barrel” Disassembly and IFT88 dispersion.** Representative timelapse images reflecting the disassembly of the tubulin barrel, as tracked via live cell imaging at 40x magnification via confocal microscopy (z-stacks, every 2.5min). RABL2-LAP (Green) + IFT88-mCherry (yellow) cells were synchronized via STAMP and stained for SiR-Tubulin (magenta). Scale bar = 20μm.

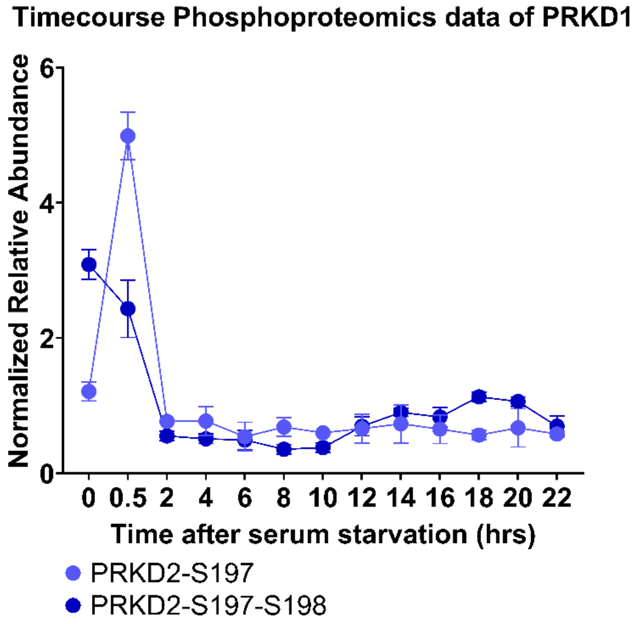


**Fig. S8. Abundance of identified phosphorylation sites on PRKD2.** Plot of the intensities of monophosphorylated [S197] and doubly phosphorylated [S197, S198] over time based on STAMP phosphoproteomics of WT RPE. Error bars = SD.

**Data S1. (separate file)**

Collision Activated Dissociation (CAD) MS2 Scans of each plotted individual phosphorylation sites discussed in the paper.
